# Lamins and lineage-relevant transcription factors coordinate gene expression in lineage development

**DOI:** 10.64898/2026.04.30.722071

**Authors:** Sara Debic, Xiaobin Zheng, Jiabiao Hu, Lidya Kristiani, Reni Marsela, Youngjo Kim, Yixian Zheng

## Abstract

**Highlights:**

- Lamin-A and lamin-B1 are essential for midgestational embryogenesis.
- Lamin-A/B1 are required for proper yolk sac endoderm (YSE) gene regulation.
- Lamin-A/B1 maintain LADs organization and chromatin interactions in YSE.
- Lamin-A/B1 and YSE transcription factors support proper YSE gene expression.

Lamins are intermediate filament proteins functioning as ubiquitous structural components of the nuclear lamina that interact with and organize the <u>L</u>amina-<u>A</u>ssociated chromatin <u>D</u>omains (LADs). LADs remodel during development and lamins maintain LADs and gene expression profile specific to a given cell type. How ubiquitous lamins achieve cell-type-specific functions during development remains unknown. We show lamin-A and -B1 are required for mouse midgestational embryogenesis and maintain LADs, 3D chromatin interactions, and gene expression in the yolk sac endoderm (YSE). Both lamin-regulated genes and remodeled LADs in YSE cells contain binding motifs of YSE-relevant transcription factors. By analyzing changes in chromatin interactions upon lamin-A and -B1 knockout, we reveal that chromatin neighborhoods maintained by these lamins can influence gene expression orchestrated by YSE-relevant transcription factors. Our findings explain how the ubiquitously expressed lamins can collaborate with lineage-relevant transcription factors to maintain LADs and gene expression programs in specific cell types.

## Introduction

Organismal development relies on highly orchestrated transcriptional programs. These transcriptional programs are intimately linked to the hierarchical three-dimensional (3D) organization of the genome, which provides a structural framework that both enables and reinforces gene expression^1^. At a scale of 100 kilobases to one million base pairs, the mammalian genome is folded into topologically-associating domains (TADs) that can constrain enhancer–promoter communication^2–4^. At a larger scale, the genome can be partitioned into two compartments based on chromatin interactions, with interactions occurring more frequently within a compartment than between the two^5^. These compartments correspond to transcriptionally active euchromatin and transcriptionally inactive heterochromatin, and are arbitrarily assigned as the A compartment and B compartment, respectively^5^. A significant fraction of B compartment heterochromatin lies underneath the nuclear lamina^6–8^, a proteinaceous layer lining the inner nuclear membrane^9^. These chromatin regions are known as the nuclear <u>L</u>amina-<u>A</u>ssociated chromatin <u>D</u>omains (LAD)^6^. LADs vary across cell types, and these differences are thought to contribute to cell-type-specific gene expression programs^10–12^. Despite significant progress in mapping LADs in different cell types^10,13–15^, how LAD organization influences transcriptional programs in a given cell type during development remains unclear.

One challenge in understanding how LAD organization influences cell-type-specific gene expression is that nuclear lamina proteins are often found in a large number of cell types^9^. Of the ∼100 reported proteins that form the nuclear lamina, the lamin intermediate filament proteins serve as the major structural component^9^. Lamins assemble into a meshwork beneath the inner nuclear envelope that interacts with other chromatin binding proteins, nuclear lamina proteins, inner nuclear envelope proteins, and the nucleoplasmic face of both Linker of Nucleoskeleton and Cytoskeleton (LINC) complexes and Nuclear Pore Complexes (NPCs)^16^. Mammals such as mice contain three lamin genes, *Lmna*, *Lmnb1*, and *Lmnb2*, which encode lamin-A and -C (abbreviated as lamin-A in this manuscript), lamin-B1, and lamin-B2, respectively^16^. During mouse development, B-type lamins are expressed in all cell types, while lamin-A is expressed in a temporally restricted manner, with expression onset depending on the cell type^17,18^. Despite their widespread expression, studies have shown that knockout of lamin-A or -B1 in various cell types results in transcriptional downregulation of genes important for the specific function of a given cell type, while genes associated with the functions of many other cell types are upregulated^19–22^. How the ubiquitously expressed lamins maintain the expression of genes relevant for cell-type-specific functions while at the same time repressing off-lineage and broadly expressed (broad-lineage) genes has remained unknown.

Studies in mouse embryonic stem cells (mESCs) have shown that lamins influence gene expression by organizing LADs to maintain 3D chromatin interactions^23^. This is further supported by later findings in other cell types derived from different cell lineages^21,24–27^. To understand how lamins facilitate specific gene expression profiles by organizing LADs in a given cell type during development, it is critical to investigate how its LADs are remodeled during differentiation. Understanding developmental LAD reorganization in this manner may shed light on how lamin knockout alters cell-type-specific LAD architecture and 3D chromatin interactions, and whether these changes may impact cell-type-specific gene expression.

Previous studies have shown that whole-body single-knockout (KO) of lamin-A results in postnatal lethality^28–30^, while single- or double-KO of lamin-B1 and lamin-B2 results in peri-natal death^31,32^. These findings serve the basis of the prevailing view that lamins are not essential for embryogenesis, and as such few efforts have examined how lamins maintain cell-type-specific transcriptional programs during development. While lamin-A is expressed at low levels in mESCs^33^, its expression is clearly evident in some cell types during embryogenesis, along with B-type lamins^17,18^. It is therefore important to explore whether lamin-A could act redundantly with B-type lamins to support embryogenesis. One challenge in studying lamins’ role in genome organization and gene regulation is that lamin knockout can cause DNA damage and apoptosis, depending on the cell type and which lamin is removed^31,32,34–36^. Therefore, to understand the role of lamins in genome organization and transcription during development without confounding effects caused by DNA damage, it is important to identify cell types that do not suffer from DNA damage upon lamin knockout.

By thoroughly analyzing the potential redundant roles of all three lamins in mouse development, we discover that lamin-A and -B1 are required for midgestational embryogenesis. Since the yolk sac endoderm (YSE) cells do not incur DNA damage upon lamin-A and -B loss, we are able to demonstrate a role for lamins in supporting YSE cell LAD organization, which in turn maintains expression of genes relevant to YSE cell functions while repressing off-lineage and broad-lineage gene expression. Our findings shed light on how the ubiquitously expressed lamins maintain developmental gene expression programs by coordinating with YSE-relevant transcription factors.

## Results

### Lamin-A and -B1 loss disrupts gene expression in YSE cells and contributes to midgestational embryonic lethality

To investigate if lamins serve functions during embryogenesis, we created male mice that are heterozygous for all three lamin genes and contain homozygous Cre recombinase alleles driven by the beta-actin promoter (*Actb-Cre^+/+^; Lmna^+/Δ^Lmnb1^+/-^Lmnb2^+/-^*). We also created female mice containing homozygous floxed alleles of all three lamin genes (*Lmna^f/f^Lmnb1^f/f^Lmnb2^f/f^*). Intercrossing of these mice showed that lamin triple-knockout (TKO) embryos and embryos lacking lamin-A and lamin-B1 (lamin-A/B1) but heterozygous for lamin-B2 (*Lmna^Δ/Δ^Lmnb1^-/-^Lmnb2^+/-^*) displayed similar growth defects compared to the control triple heterozygote (*Lmna^+/Δ^Lmnb1^+/-^Lmnb2^+/-^*) littermates at embryonic day (E)9.5 (Figure S1A). No viable embryos of either genotype were recovered at E11.5 (data not shown). This shows that lamin-A and lamin-B1, but not lamin-B2, are important for supporting midgestational embryo development.

Since the lamin-A/B1 double-knockout (DKO) embryos had one copy of the lamin-B2 gene, we next analyzed lamin-A/B1 DKO embryos with two copies of the lamin-B2 gene by crossing *Actb-Cre^+/+^; Lmna^+/Δ^Lmnb1^+/-^*male mice with *Lmna^f/f^Lmnb1^f/f^* female mice. We examined embryos from E7.5 to E9.5 and observed that lamin-A/B1 DKO (*Lmna^Δ/Δ^Lmnb1^-/-^Lmnb2^+/+^*) embryos exhibited a similar growth defect as lamin TKO embryos (Figure 1A). A noticeable growth defect in lamin-A/B1 DKO embryos started at E8.5 compared to lamin-A/B1 double heterozygote (*Lmna^+/Δ^Lmnb1^+/-^*) littermate controls (Figure 1A). The growth defect continued and some embryos failed to turn by E9.5 (Figure 1A). By E11.5, we could not retrieve any viable lamin-A/B1 DKO embryos (Figure 1B). These studies show that lamin-A/B1 DKO causes significantly earlier lethality than the postnatal lethality of lamin-A single-KO or the perinatal lethality of lamin-B1 single-KO, lamin-B2 single-KO, or DKO of lamin-B1/B2^28–32^. Thus, lamin-A and -B1 play essential roles during midgestational mouse development.

**Figure 1.**
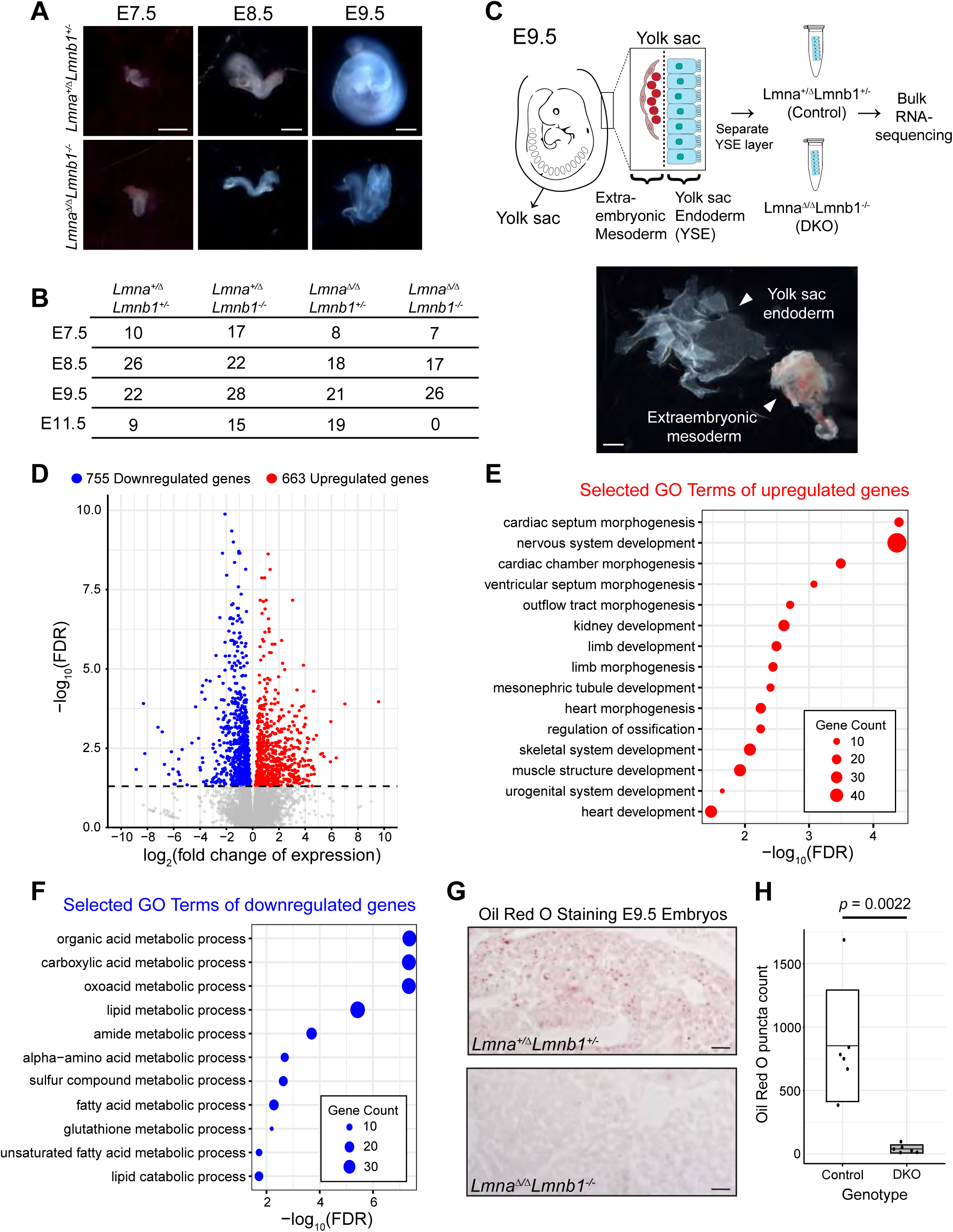
Lamin-A and -B1 maintain transcriptional programs in yolk sac endoderm (YSE) cells and are required for midgestational embryo development. (A) Representative images of control (*Lmna^+/Δ^Lmnb1^+/-^*) and lamin-A/B1 double-knockout (DKO; *Lmna^Δ/Δ^Lmnb1^-/-^*) embryos at embryonic day (E)7.5, E8.5, and E9.5. Scale bar, 500 µm. (B) Numbers of embryos of each genotype recovered from crosses of *Actb-Cre^+/+^*; *Lmna^+/Δ^Lmnb1^+/-^* and *Lmna*^f/f^*Lmnb1^f/f^* mice from E7.5 to E11.5. No lamin-A/B1 DKO embryos were recovered at E11.5 (Chi-squared test: χ^2^ = 19.05, degrees of freedom = 3, *p* = 0.00027). (C) Schematic of yolk sac endoderm (YSE) cell isolation and bulk RNA-seq workflow. Micrograph shows an isolated YSE cell layer. Scale bar, 500 µm. (D) Volcano plot of differentially expressed genes in lamin-A/B1 DKO versus control YSE cells identified by bulk RNA-seq. Downregulated and upregulated genes with a false discovery rate (FDR) < 0.05 are highlighted in blue and red, respectively. (E and F) Selected Gene Ontology (GO) terms significantly enriched among genes upregulated (E) or downregulated (F) following lamin-A/B1 loss in YSE cells. (G) Representative Oil Red O staining of magnified regions from control and lamin-A/B1 DKO embryonic sections. Scale bar, 10 µm. (H) Quantification of Oil Red O puncta per section from control and lamin-A/B1 DKO embryos. Boxes indicate the mean ± SD; points represent individual sections. N = 2 biological replicates per genotype; *p* = 0.006, Student’s t-test.

To further understand the role of lamin-A/B1 during embryogenesis, we next examined cell types that express lamin-A and -B1 at E8.5 when the growth defect of lamin-A/B1 DKO embryos became apparent. While lamin-B1 and -B2 are expressed in all tissues throughout mouse development^17^, lamin-A is not strongly expressed in the embryo proper at E8.5^18^. Instead, lamin-A-positive cells are found in the extraembryonic yolk sac and trophoblast tissue at this developmental stage^17^. Our immunofluorescence staining analyses reveal that all three lamins are present in the yolk sac endoderm (YSE) cells of control mice, while lamin-A/B1 are absent in the lamin-A/B1 DKO cells (Figure S1B-S1D). We observed a noticeable increase in lamin-B2 immunofluorescence in YSE cells upon lamin-A/B1 DKO (Figure S1D), a compensation phenomenon observed upon deletion of members of a gene family^37,38^. Nonetheless, since the yolk sac supports mouse embryo growth by supplying nutrients^39–41^ before the placenta starts transporting nutrients to the embryo at ∼E10^42^, the apparent growth defect at E8.5 of lamin-A/B1 DKO embryos reveal a role for lamin-A/B1 in the yolk sac to support embryo growth before the placenta becomes functional.

Studies in different cell types have shown that lamin deletion can cause DNA damage or apoptosis^31,32,34–36^, as well as altered distribution of nuclear peripheral proteins such as the NPCs^43,44^, all of which could contribute to YSE cell dysfunction. We used immunofluorescence staining to validate antibodies detecting DNA damage and apoptosis in tissues that are known to suffer from DNA damage and apoptosis (data not shown). Using these validated antibodies, we found weak DNA damage and apoptosis signals that are similar in both control and lamin-A/B1 DKO YSE cells (Figure S1E-S1H). We also did not observe defects in nuclear pore distribution in the lamin-A/B1 DKO YSE cells at E9.5 (Figure S1I). Therefore, YSE cells offer an opportunity to investigate the direct role of lamins in regulating genome organization and transcription during development.

To investigate if lamin-A/B1 DKO alters transcriptional programs, we isolated YSE cells by enzymatically disrupting the extracellular matrix between the YSE cell epithelial layer and the extraembryonic mesoderm. By carefully peeling the YSE cell epithelial layer off the mesoderm, we were able to isolate the YSE cells with more than 90% purity (Figure 1C)^45^. Bulk RNA sequencing (RNA-seq) on the isolated E9.5 YSE cells from lamin-A/B1 DKO and control embryos identified 663 upregulated and 755 downregulated genes (False Discovery Rate < 0.05) upon lamin-A/B1 loss (Figure 1D). Gene Ontology (GO) term analyses showed that the upregulated genes are involved in the functions of alternate cell lineages, such as the nervous system, heart, and kidneys (Figure 1E; Table S1), and include genes broadly expressed in many lineages (see below). This is consistent with previous reports that lamins suppress the expression of off- or broad-lineage genes^19–22^. For example, we observed upregulation of several off-lineage genes, including the *Hoxb1* gene involved in hindbrain and cranial nerve development (Table S1)^46,47^. We also found upregulated genes involved in developmental signaling pathways broadly expressed in many cell types (broad-lineage genes), including genes encoding members of the Wnt, Notch, Hedgehog, and FGF signaling pathways that are expressed at low levels in control YSE cells (Table S1).

GO term analyses of genes downregulated upon lamin-A/B1 loss in YSE cells revealed many of them encode proteins involved in lipid metabolism, a core function of YSE cells (Figure 1F; Table S1)^39–41^. For example, we found downregulation of the *Apoa2* gene, encoding Apolipoprotein A2, a structural component of high-density lipoproteins that transport cholesterol and other lipids in the bloodstream (Table S1)^48^. Consistent with the transcriptional downregulation of lipid transport genes, Oil Red O staining of histological sections of E9.5 embryos revealed a significant reduction of neutral lipids upon lamin-A/B1 DKO as compared to control littermates (Figure 1G, 1H). Thus, lamin-A and -B1 contribute to midgestational mouse development in part by maintaining the expression of genes required for YSE cell-mediated lipid transport to the embryo, while repressing the expression of off-lineage and broad-lineage genes.

### Binding motifs of YSE-relevant transcription factors are enriched in lamin-regulated genes

To understand how lamin-A/B1 maintain gene expression and repression important for YSE cell function, we investigated whether transcription factors with roles in YSE cells could be involved. We explored binding motifs of transcription factors enriched in the dysregulated genes upon lamin-A/B1 loss. Since accessible chromatin regions often contain enhancers and promoters that are bound by transcription factors^49^, we performed the <u>A</u>ssay for <u>T</u>ransposase <u>A</u>ccessible <u>C</u>hromatin using <u>seq</u>uencing (ATAC-seq) in wild-type E9.5 YSE cells and identified accessible chromatin regions using peak calling (Figure 2A, S2A, S2B, see Methods). We then used Homer to assign ATAC-seq peaks to genes and further identified those peaks associated with the lamin-A/B1-regulated genes (Figure S2C). Strikingly, we found that the ATAC-seq peaks associated with both downregulated and upregulated genes are significantly enriched for binding motifs of Hepatocyte Nuclear Factor 1-beta (HNF1ϕ3), HNF4α, GATA-binding protein 4 (GATA4), and GATA6, which are transcription factors with well-established roles in YSE development (Figure 2B-2E; Table S2)^50–53^.

**Figure 2.**
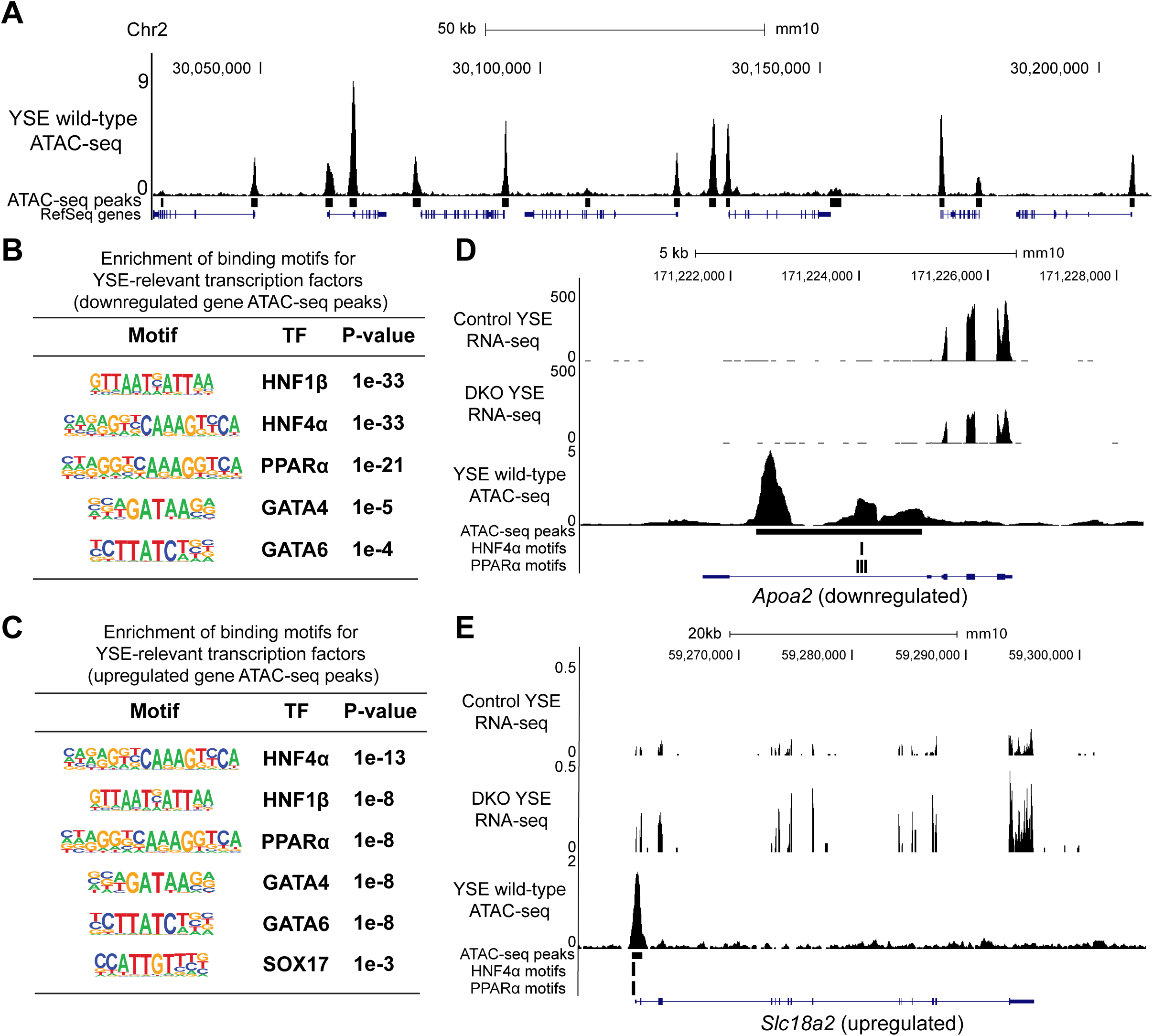
Binding motifs of YSE-relevant transcription factors are associated with genes dysregulated upon lamin-A/B1 loss. (A) Representative genome browser track of ATAC-seq signal from YSE cells. ATAC-seq peaks were called using MACS2 and ATAC-seq tracks are shown as reads normalized to 1 million mapped reads. (B and C) Significant enrichment of binding motifs for YSE-relevant transcription factors in ATAC-seq peaks associated with downregulated (B) or upregulated (C) genes following lamin-A/B1 loss. (D and E) Genome browser tracks showing a representative downregulated gene *Apoa2* (D) and a representative upregulated gene *Slc18a2* (E) with associated ATAC-seq peaks containing binding motifs for the indicated YSE-relevant transcription factors. RNA-seq tracks are shown as reads normalized to total library size, and ATAC-seq tracks are shown as reads normalized to 1 million mapped reads.

Additionally, we found that both the downregulated and upregulated genes are highly enriched for binding motifs of Peroxisome Proliferator-Activated Receptor alpha (PPARα) (Figure 2B-2E; Table S2), a transcription factor with known roles in regulating gene expression in lipid-handling tissues^54^. Interestingly, the upregulated genes also display strong enrichment for binding motifs of several SOX family transcription factors (Table S2). These include SOX proteins with roles in alternate lineage development such as SOX3^55^, and SOX17, a transcription factor involved in YSE development (Figure 2C; Table S2)^56^. These analyses suggest lamin-A/B1 could collaborate with YSE-relevant transcription factors to influence both gene expression and repression.

### Binding motifs of YSE-relevant transcription factors are enriched in chromatin regions undergoing LAD remodeling in YSE cells during development

We next explored whether YSE-relevant transcription factors could contribute to YSE cell LAD remodeling during development. We mapped LADs in wild-type E9.5 YSE cells using <u>C</u>leavage <u>U</u>nder <u>T</u>argets and <u>R</u>elease <u>U</u>sing <u>N</u>uclease (CUT&RUN) with antibodies targeting lamin-A, -B1, or -B2. Random IgG was used as a control for non-specific binding. LAD profiles from the three lamin isoforms were highly concordant (Pearson correlations > 0.85; Figure S3A, S3B), so we averaged z-scores at 5-kb resolution for downstream analyses and LAD calling. Since previous studies have shown LAD remodeling is accompanied by epigenetic changes during cell differentiation^57^, we next used CUT&RUN to map H3K9me2, H3K9me3, H3K27me3, H3K27ac, and H3K4me3 in wild-type E9.5 YSE cells. To analyze LADs and their epigenome features in YSE cells, we used the <u>Hi</u>stone and lamina <u>Lands</u>capes (HiLands) model developed for mESCs to annotate YSE cell chromatin states based on both LADs and epigenetic information^58^. By combining the YSE cell epigenome maps with ATAC-seq and LAD mapping, we used a hidden-Markov model (HMM) to define six HiLands in YSE cells as HiLands-Purple (P), -Blue (B), - Green (G), -Yellow (Y), -Orange (O), and -Red (R) (Figure 3A, 3B).

**Figure 3.**
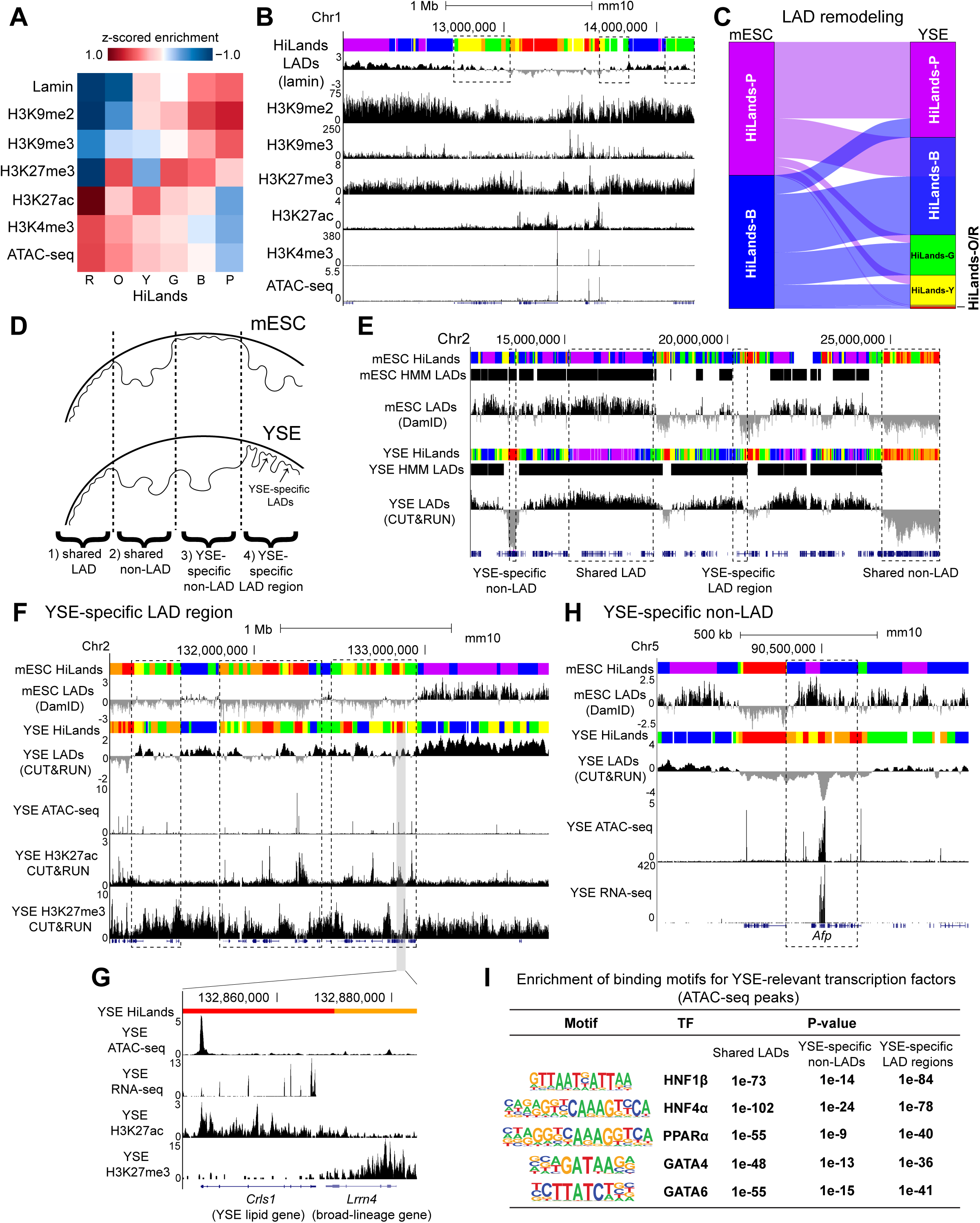
LAD remodeling between YSE cells and mESCs. (A) Heatmap showing averaged z-scored enrichment of CUT&RUN reads from all three lamin isoforms (lamin), histone modification CUT&RUN (H3K9me2, H3K9me3, H3K27me3, H3K27ac, H3K4me3), and ATAC-seq signal across the six HiLands chromatin states of YSE cells. Lamin enrichment was calculated as fold-change over IgG CUT&RUN prior to z-scoring; histone modification CUT&RUN and ATAC-seq enrichment was z-scored directly. (B) Representative genomic region showing the six HiLands in wild-type YSE cells with genome browser tracks of LADs determined by lamin CUT&RUN enrichment compared to IgG CUT&RUN, histone modification CUT&RUN, and ATAC-seq. Boxed regions highlight fragmented LADs characterized by HiLands-G and -Y. (C) Alluvial plot showing transitions of HiLands-P and HiLands-B LADs in mESCs to their corresponding HiLands in YSE cells. (D) Schematic illustration of the four categories of LADs and non-LADs in mESCs and YSE cells. (E) Representative genomic region illustrating the four categories of LADs and non-LADs in mESCs and YSE cells. Genome browser tracks include published mESC HiLands and LADs mapped by lamin-B1 Dam-ID, as well as hidden-Markov model (HMM)-called mESC LAD annotations generated in this study. Additional tracks include YSE HiLands, YSE HMM-called LAD annotations, and YSE LADs mapped by lamin CUT&RUN all generated in this study. Dashed boxes indicate examples of YSE-specific non-LADs, shared LADs, YSE-specific LAD regions, and shared non-LADs. (F) Representative fragmented YSE-specific LAD regions (dashed boxes). Genome browser tracks include published mESC LADs mapped by lamin-B1 Dam-ID and HiLands, along with YSE HiLands, LADs mapped by lamin CUT&RUN, ATAC-seq, H3K27ac CUT&RUN, and H3K27me3 CUT&RUN generated in this study. (G) Zoomed-in view of the gray highlighted region in (F) showing an example expressed YSE gene involved in lipid metabolism within HiLands-R and a repressed broad-lineage gene within HiLands-O regions that interrupt YSE-specific LADs. Genome browser tracks include YSE HiLands, ATAC-seq, RNA-seq, as well as H3K27ac and H3K27me3 CUT&RUN generated in this study. (H) Representative YSE-specific non-LAD (dashed boxed) containing the highly expressed YSE gene *Afp*. Genome browser tracks include published mESC LADs mapped by lamin-B1 Dam-ID and HiLands, along with YSE HiLands, LADs mapped by lamin CUT&RUN, ATAC-seq, and RNA-seq generated in this study. (I) Enrichment of binding motifs for YSE-relevant transcription factors in YSE ATAC-seq peaks associated with shared LADs, YSE-specific non-LADs, and YSE-specific LAD regions.

Similar to mESCs, YSE cells exhibit the lowest gene expression in HiLands-P followed by HiLands-B, whereas the highest gene expression is found in HiLands-R (Figure S3C). While LADs in mESCs mainly correspond to HiLands-P and -B^58^, LADs in YSE cells are distributed among HiLands-P, -B, -G, and -Y (Figure 3A, 3B, S3D). The LADs in mESCs^58^ and YSE cells represented by HiLands-P and -B share similar epigenome features, with the HiLands-P LADs enriched for heterochromatin modifications (H3K9me2 and/or H3K9me3) and the HiLands-B LADs enriched for H3K27me3 (see Figure 3A for YSE cells). Interestingly, the YSE cell LADs characterized by HiLands-G and -Y are often fragmented and found at the end of HiLands-B LADs (see dashed boxes in Figure 3B). Lastly, HiLands-O and HiLands-R in both mESCs^58^ and YSE cells are found mainly in the nuclear interior as non-LAD regions (see Figure S3D for YSE cells). To investigate how the LADs in YSE cells are remodeled from their progenitor cells during development, it is informative to compare YSE cell LADs to those in the inner cell mass (ICM) cells because ICM cells differentiate into the YSE lineage^59^. Since the LADs mapped in mESCs derived from ICM cells closely resemble the ICM cell LADs mapped in mouse preimplantation blastocysts^60^, we used the HMM-called LADs and HiLands modeling from mESCs for the comparison. We find that the vast majority of mESC LADs are characterized by HiLands-P and - B, and they are remodeled to become different HiLands in YSE cells (Figure 3C). For HiLands-P LADs in mESCs, ∼57% remain HiLands-P LADs, ∼29% become HiLands-B LADs, ∼13% become HiLands-G or -Y, and only ∼1% become HiLands-O or -R in YSE cells (Figure 3C). For HiLands-B LADs in mESCs, ∼44% remain HiLands-B, ∼14% become HiLands-P, ∼40% become HiLands-G or -Y, and ∼2% become HiLands-O and -R in YSE cells (Figure 3C).

Comparing LADs and non-LADs between mESCs and YSE cells revealed four types of chromatin regions (Figure 3D). As expected, the mESCs and YSE cells have large regions of 1) shared LADs and 2) shared non-LADs, which represent the majority of the genome (Figure 3D, 3E, S3E, S3F). The 3) YSE-specific non-LADs are chromatin regions that are LADs only in mESCs (Figure 3D, 3E, S3E). The YSE-specific LADs refer to LADs present only in YSE cells (Figure 3D, 3E, S3F). Since the YSE-specific LADs are frequently interrupted by short non-LAD regions (Figure 3D, see dashed boxes in Figure 3F), we refer to these regions as 4) YSE-specific LAD regions.

To investigate whether lineage-relevant transcription factors could be involved in LAD remodeling, we analyzed ATAC-seq peaks in the shared LADs, YSE-specific LAD regions, YSE-specific non-LADs, and shared non-LADs in YSE cells. We found that the shared LAD regions contain infrequent ATAC-seq peaks in YSE cells, which coincide with local reductions in nuclear lamina association (Figure S3G, S3H). The YSE-specific LAD regions also contain ATAC-seq peaks, which often coincide with chromatin regions that are non-LADs including HiLands-O and -R (Figure 3F). The HiLands-R regions interrupting YSE-specific LADs include genes expressed in YSE cells that display strong ATAC-seq peaks (Figure 3G). By contrast, the HiLands-O regions interrupting YSE-specific LADs contain H3K27me3-repressed developmental genes exhibiting weak ATAC-seq peaks (Figure 3G), which is consistent with previous reports showing ATAC-seq signal can colocalize with repressive histone modifications^61^. As expected, the YSE-specific non-LADs contain a high density of ATAC-seq peaks, especially at genes that are highly expressed and have known functions in YSE cells such as *Afp* (Figure 3H). Lastly, the shared non-LAD regions in YSE cells contain the majority of ATAC-seq peaks and actively expressed genes, as expected (Figure S3I, S3J).

To investigate potential factors involved in LAD remodeling in YSE cells, we next performed motif enrichment analyses of ATAC-seq peaks found in shared LAD and YSE-specific LAD regions, as well as the YSE-specific non-LADs. Strikingly, ATAC-seq peaks across all these regions are significantly enriched for binding motifs of YSE-relevant transcription factors, including HNF1ϕ3, HNF4α, GATA4, GATA6, and PPARα (Figure 3I). To explore whether the YSE-specific LADs represented by HiLands-B, G, and -Y contain specific binding motifs for factors that may aid LAD formation, we also analyzed motif enrichment in these regions. We found enrichment for motifs of several transcription factors expressed in YSE cells with known repressive functions, including PBX2 and FOXO1 (Figure S3K; Table S3)^62,63^, suggesting that transcriptional repressors may facilitate YSE-specific LAD formation. Thus, YSE-related transcription factors may facilitate LAD remodeling in YSE cells during lineage development.

### Lamin-A and -B1 support LAD organization and 3D chromatin interactions in YSE cells

Lamins are known to influence gene expression by organizing LADs^21,23–27^. To explore whether lamin-A/B1 may collaborate with YSE-relevant transcription factors to support gene expression, it is important to understand how lamin-A/B1 loss changes LAD organization and 3D chromatin interactions and how such changes may impact these transcription factors in YSE cells. We attempted to map LADs in the E9.5 lamin-A/B1 DKO YSE cells but found that the smaller lamin-A/B1 DKO yolk sacs yielded lower numbers of YSE cells compared to controls, and we failed to map LADs using the CUT&RUN method. However, since LADs can be inferred from 3D chromatin interactions^7,8^, we mapped 3D chromatin interactions in E9.5 control and lamin-A/B1 DKO YSE cells using a <u>Hi</u>gh-throughput chromosome conformation <u>C</u>apture (Hi-C) protocol developed for low cell numbers. This approach yielded high-quality Hi-C libraries in multiple biological replicates of control and lamin-A/B1 DKO YSE cells (Table S4).

The Hi-C contact matrices were first normalized using the Knight-Ruiz algorithm to correct for systematic biases (see Methods). By analyzing contact frequency decay curves, which plot the average chromatin interactions against the linear genomic distance between interacting sites, we found the Hi-C replicates displayed good consistency (Figure S4A). Thus, we merged unique reads from individual Hi-C replicates corresponding to each genotype for all downstream analyses. Visualization of chromatin interactions as heatmaps in control and lamin-A/B1 DKO YSE cells revealed characteristic TAD structures (Figure 4A, 4B). To directly compare the TADs between control and lamin-A/B1 DKO YSE cells, we called TAD boundaries using the insulation score method. This method quantifies chromatin interactions within sliding genomic windows, and windows displaying local contact frequency minima are assigned as TAD boundaries. This analysis revealed overall similar insulation scores and TAD boundaries between control and lamin-A/B1 DKO YSE cells (Figure 4A, 4B, S4B, S4C). Therefore, the loss of lamin-A/B1 does not substantially alter TAD organization in YSE cells, similar to observations made in different tissue culture cells upon knockout of various lamins^23,25^.

**Figure 4.**
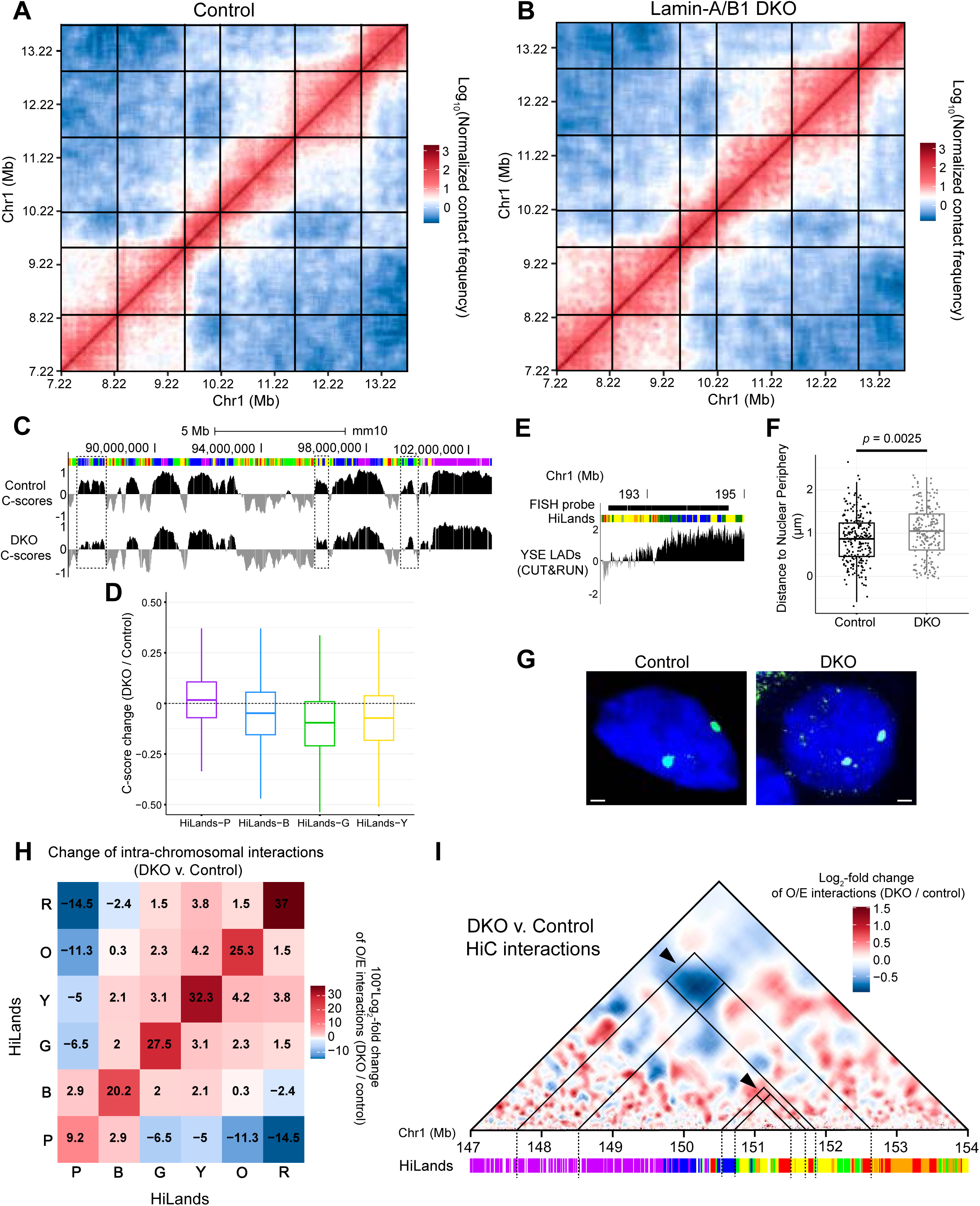
Lamin-A/B1 maintain LAD organization and 3D chromatin interactions in YSE cells. (A and B) Heatmaps showing observed Hi-C interaction frequencies at 20 kb resolution for a representative region of chromosome 1 in control (A) and lamin-A/B1 DKO (B) YSE cells. Black lines indicate TAD boundaries called using the insulation score method. Contact matrices were normalized using the Knight-Ruiz method and by total library size. (C) Representative region showing Hi-C compartment scores (C-scores) in control and lamin-A/B1 DKO YSE cells alongside YSE HiLands. Dashed boxes highlight decreased C-scores in compartment B regions characterized by HiLands-B, HiLands-G, and HiLands-Y following lamin-A/B1 loss. (D) Genome-wide quantification of C-score changes following lamin-A/B1 loss for LADs characterized by HiLands-P, HiLands-B, HiLands-G, and HiLands-Y. Boxes show the interquartile range, center lines indicate medians, and whiskers extend to 1.5 times the interquartile range. (E) Genome browser track of LADs mapped by lamin CUT&RUN in YSE cells showing the FISH probed LAD region encompassing HiLands-B, HiLands-G, and HiLands-Y. (F) Quantification of the distance from the FISH probe to the nuclear periphery (DAPI) in control and lamin-A/B1 DKO YSE cells. Boxes indicate interquartile range, center lines indicate medians, and whiskers extend to 1.5 times the interquartile range. N = 2 biological replicates; 225 control and 198 lamin-A/B1 DKO nuclei analyzed. *p* = 0.0025, Wilcoxon rank-sum test. (G) Representative confocal images of control and lamin-A/B1 DKO YSE cells showing the FISH probed region (green). Single z-planes are shown. Scale bar, 2 µm. (H) Heatmap showing genome-wide fold-changes of observed/expected (O/E) intra-chromosomal Hi-C interactions within HiLands and between different HiLands pairs upon lamin-A/B1 loss. (I) Heatmap of a representative chromosome 1 region showing fold-changes in observed/expected (O/E) Hi-C interactions between control and lamin-A/B1 DKO YSE cells. A YSE HiLands track is included; boxed regions highlight reduced interactions between HiLands-P and other HiLands, and increased interactions between HiLands-B and other HiLands.

To infer LADs, we performed compartment analyses based on Hi-C. Previous studies showed that the genome can be separated into A and B compartments^5^, with the B compartment containing mostly heterochromatic LADs and the A compartment containing nuclear interior chromatin^7,8^. The A and B compartments are modeled based on their 3D chromatin interactions which are higher within each compartment than between the two^5^. We mapped A and B compartments in the E9.5 control and lamin-A/B1 DKO YSE cells using the CscoreTool developed in our lab^64^. Calculations were performed using our control and lamin-A/B1 DKO YSE Hi-C datasets at 20 kb resolution. The C-scores represent the probability of individual chromatin regions being in the A or B compartments. We assigned the positive or negative C-scores to correspond to the B and A compartments, respectively (Figure 4C). As expected, the B compartment scores in control YSE cells display strong correlation with YSE LAD mapping data (Pearson correlation = 0.77; Figure S4D, S4E). Accordingly, the repressive HiLands-P and -B LADs have mainly positive C-scores, HiLands-G and -Y display both positive and negative C-scores, while the interior HiLands-O and -R in the A compartment have mainly negative C-scores (Figure 4C, S4F).

Upon lamin-A/B1 loss, we observed LADs characterized by HiLands-B, -G, and -Y exhibit a decrease in C-scores genome-wide, indicating their reduced compartment B association (Figure 4C, 4D). By contrast, HiLands-P LADs did not exhibit a similar decrease in C-scores genome-wide, indicating that HiLands-P LADs largely retain compartment B identity upon lamin-A/B1 loss (Figure 4C, 4D). The reduction in C-scores for LADs characterized by HiLands-B, -G, and - Y suggests detachment of these regions from the nuclear periphery upon lamin-A/B1 loss. This movement into the nuclear interior may increase their interactions with interior chromatin in compartment A and reduce their interactions with chromatin in compartment B. To experimentally verify that lamin-A/B1 loss causes detachment of LADs characterized by HiLands-B, -G, and -Y from the nuclear periphery, we performed fluorescent in situ hybridization (FISH) using probes corresponding to a LAD region composed of mainly HiLands-B, -G, and -Y (Figure 4E). This probed region also displays decreased C-scores upon lamin-A/B1 loss (Figure S4G). FISH analyses showed a significant re-localization of this probed region away from the nuclear periphery in lamin-A/B1 DKO YSE cells compared to controls (Figure 4F, 4G). These analyses show that lamin-A/B1 are required to maintain LAD regions characterized by HiLands-B, -G, and -Y in YSE cells.

We next examined global chromatin interaction changes upon lamin-A/B1 loss in the context of HiLands, because the epigenetic information offered could shed light on how lamin-A/B1 influence gene expression by maintaining interactions among chromatin regions with different epigenetic features. We analyzed the change in intra-chromosomal Hi-C interactions within and between various HiLands upon lamin-A/B1 DKO. To normalize for distance-dependent effects, we calculated the observed/expected Hi-C interaction ratios for each genotype and then computed fold-changes of these ratios between lamin-A/B1 DKO and control YSE cells (see Methods). We found that HiLands-P display a genome-wide decrease in interactions with all other HiLands except for HiLands-B (Figure 4H, 4I). By contrast, HiLands-B display increased interactions with HiLands-P, -G, -Y, and -O, but decreased interactions with transcriptionally active HiLands-R (Figure 4H, 4I). The HiLands-G, -Y, -O, and -R all show increased interactions with each other (Figure 4H, 4I). These results are consistent with our compartment analysis showing that upon lamin-A/B1 loss, HiLands-B, -G, and -Y LADs, but not HiLands-P LADs, shift from compartment B to compartment A, thereby gaining interactions with HiLands in the A compartment. Additionally, we found that all the HiLands exhibit increased interactions with themselves, with the transcriptionally active HiLands-R and -Y showing the greatest increase (Figure 4H). Our findings show that lamin-A/B1 organize LADs and maintain interactions among chromatin regions with distinct epigenetic features.

### Lamin-A and -B1 maintain chromatin neighborhoods that can regulate gene expression in YSE cells

The observed detachment of LADs characterized by HiLands-B, -G, and -Y chromatin from the nuclear lamina upon lamin-A/B1 loss (Figure 4C-4G) could disrupt 3D genome interactions and local chromatin neighborhoods, thereby impacting gene expression^65–67^. To explore this, we first mapped promoters of genes dysregulated upon lamin-A/B1 loss throughout the genome, categorizing them in shared non-LADs, YSE-specific LAD regions, shared LADs, or YSE-specific non-LADs. We find ∼65% of the dysregulated gene promoters are in shared non-LADs, ∼26% in YSE-specific LAD regions, ∼9% in shared LADs, and only a few genes are in YSE-specific non-LADs (Figure 5A). We then analyzed the distance of each of these groups of dysregulated gene promoters to the nearest LAD regions characterized by HiLands-B, -G, and -Y. By plotting the linear genomic distance of the dysregulated gene promoters in shared non-LADs to their nearest HiLands-B, -G, and -Y LADs, we find these gene promoters are located farther away from these regions compared to all expressed YSE transcripts (Figure S5A). Thus, the transcriptional dysregulation of genes in the shared non-LADs is unlikely caused only by the detachment of LADs characterized by HiLands-B, -G, and -Y, but could instead result from additional indirect effects of lamin-A/B1 loss.

**Figure 5.**
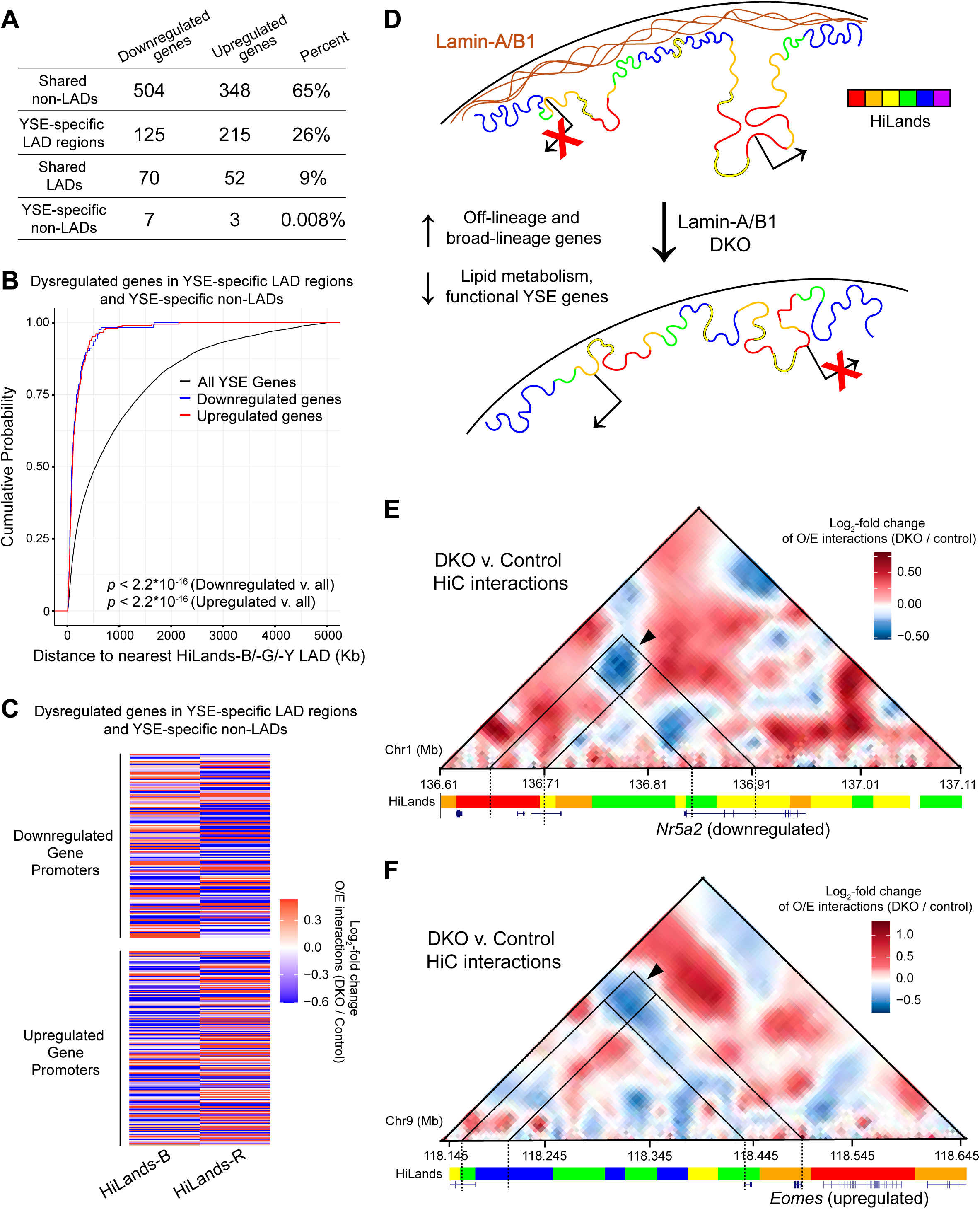
Lamin-A/B1 maintain LADs and 3D chromatin interactions to support gene expression in YSE cells. (A) Distribution of genes dysregulated upon lamin-A/B1 loss across the four categories of LADs and non-LADs in YSE cells. (B) Cumulative probability plot of distances from promoters of dysregulated genes in YSE-specific LAD regions and YSE-specific non-LADs to the nearest LAD region characterized by HiLands-B, HiLands-G, and HiLands-Y. Distances of all expressed YSE genes are shown as a control. Statistical comparisons to control were performed using the Kolmogorov-Smirnov test: upregulated vs. control, *p* < 2.2 * 10^-16^; downregulated vs. control, *p* < 2.2 * 10^-16^. (C) Heatmaps showing changes in 3D chromatin interactions between 5 kb bins covering promoters of dysregulated genes in the indicated LAD and non-LAD regions and HiLands-B and HiLands-R within ±5 megabases (Mb) from individual promoters. Downregulated genes exhibit decreased interactions with HiLands-R (*p* = 0.015, Wilcoxon rank-sum test), while upregulated genes display decreased interactions with HiLands-B (*p* = 0.020, Wilcoxon rank-sum test). Significance was assessed relative to randomized control genes (see Methods). (D) Schematic illustration of how lamin-A/B1-dependent LAD organization and chromatin interactions influence gene expression in YSE cells. Upregulated off-lineage or broad-lineage genes lose interactions with repressive HiLands-B, while downregulated lipid metabolism genes lose interactions with transcriptionally active HiLands-R. (E and F) Heatmaps showing 3D chromatin interaction changes upon lamin-A/B1 loss around the downregulated gene *Nr5a2* (E) and the upregulated gene *Eomes* (F). Boxed regions highlight decreased chromatin interactions with transcriptionally active HiLands-R (E) or repressive HiLands-B (F).

We next examined dysregulated gene promoters in shared LADs, YSE-specific non-LADs, and YSE-specific LAD regions and found they are located significantly closer to LADs characterized by HiLands-B, -G, and -Y compared to all expressed YSE transcripts (Figure 5B, S5B). This suggests detachment of LADs characterized by HiLands-B, -G, and -Y could alter the chromatin neighborhoods and transcriptional output of these groups of genes. We then analyzed how the promoters of each of these gene groups alter their chromatin interactions with surrounding HiLands upon lamin-A/B1 loss. While we did not find trends in 3D chromatin interaction changes that explain dysregulation of genes with promoters in shared LAD regions, our analyses revealed the genes with promoters in YSE-specific non-LADs and YSE-specific LAD regions exhibit trends in 3D chromatin interaction changes that help explain transcriptional dysregulation (Figure 3C). The downregulated genes in these regions show a significant relocation of promoters away from the transcriptionally active HiLands-R upon lamin-A/B1 loss compared to a randomized set of control genes (see Methods, Figure 5C; *p* = 0.015, Wilcoxon rank-sum test). By contrast, the upregulated gene promoters in YSE-specific non-LADs and YSE-specific LAD regions significantly lose interactions with repressive HiLands-B chromatin compared to a randomized set of control genes (Figure 5C; *p* = 0.020, Wilcoxon rank-sum test). Thus, decreased interactions with active HiLands-R chromatin can contribute to gene downregulation, while decreased interactions with repressive HiLands-B chromatin can contribute to gene upregulation upon lamin-A/B1 loss (Figure 5D). These analyses reveal that transcriptional changes of ∼26% of the dysregulated genes can be explained by their promoters losing interactions with transcriptionally active or inactive chromatin neighborhoods. Additionally, the downregulated and upregulated genes in this subset are enriched for lipid metabolism-related or off-/broad-lineage GO terms, respectively (Figure S5C, S5D; Table S5).

Since the observed 3D chromatin interaction changes only explain a subset of the dysregulated genes upon lamin-A/B1 loss, we next explored whether some of these genes may function as upstream regulators that in turn alter the expression of other genes in YSE cells. We indeed found several such upstream regulators (Table S5). For example, the *Nr5a2* gene is significantly downregulated upon lamin-A/B1 loss (Table S5), and its loss of 3D chromatin interactions with active HiLands-R regions may contribute to this repression (Figure 5E, S5E, S5F). *Nr5a2* encodes a nuclear hormone receptor that regulates downstream genes involved in lipid metabolism^68^. The downregulation of *Nr5a2* due to 3D chromatin interaction changes caused by lamin-A/B1 loss can result in downregulation of additional lipid metabolism genes throughout the genome. Indeed, we found ∼9% of downregulated genes contain binding motifs for Nr5a2 in their associated ATAC-seq peaks (Table S5).

We next analyzed potential upstream regulators whose upregulation due to lamin-A/B1 loss can be explained by 3D chromatin changes and identified *Eomes* (Table S5). *Eomes* is a transcription factor that regulates development of diverse lineages and influences the expression of downstream genes^69^. The region encompassing the *Eomes* locus loses interactions with repressive HiLands-B upon lamin-A/B1 loss, which correlates with its transcriptional upregulation (Figure 5F, S5G, S5H). The increase in *Eomes* expression upon lamin-A/B1 loss may cause upregulation of its target genes throughout the genome. Indeed, we found ∼40% of upregulated genes contain binding motifs for Eomes in their associated ATAC-seq peaks (Table S5). Therefore, dysregulation of upstream regulators due to altered chromatin neighborhoods can result in additional gene dysregulation independent of 3D chromatin interaction changes upon lamin-A/B1 loss in YSE cells.

### Lamin-A and -B1 maintain chromatin neighborhoods to support gene expression regulated by YSE-relevant transcription factors

To further explore how the altered chromatin neighborhoods upon lamin-A/B1 loss can cause the dysregulation of genes found in YSE-specific non-LAD and YSE-specific LAD regions, we analyzed the ATAC-seq peaks associated with these genes. We found the majority of the peaks contain binding motifs for YSE-relevant transcription factors (Figure S6A; Table S6). Since transcription factor activity can be impacted by whether the target genes are in active or repressive chromatin neighborhoods^49,65–67^, we analyzed chromatin interaction changes around these genes associated with YSE-relevant transcription factor motifs. We found the downregulated genes associated with YSE-relevant transcription factor motifs significantly lose chromatin interactions with the active HiLands-R compared to a set of randomized control genes (Figure S6B; *p* = 0.043, Wilcoxon rank-sum test). These interaction changes can bring the genes into less active or repressive chromatin neighborhoods, thereby limiting the activity of YSE-relevant transcription factors and causing gene downregulation upon lamin-A/B1 deletion. Concomitant with these chromatin interaction alterations, we observed a reduction in the active enhancer mark H3K27ac at these downregulated gene promoters (Figure 6A). For the upregulated genes associated with YSE-relevant transcription factor motifs, we found a significant loss in their chromatin interactions with repressive HiLands-B compared to a set of randomized control genes (Figure S6B; *p* = 0.013, Wilcoxon rank-sum test). These decreased interactions with repressive chromatin neighborhoods can promote activity of YSE-relevant transcription factors, thereby contributing to increased gene expression. Consistently, these upregulated genes exhibit a loss in the repressive histone mark H3K27me3 at their promoters (Figure 6B). These findings suggest that lamin-A/B1 could maintain chromatin interaction neighborhoods around YSE-relevant transcription factors to support YSE gene expression programs.

**Figure 6.**
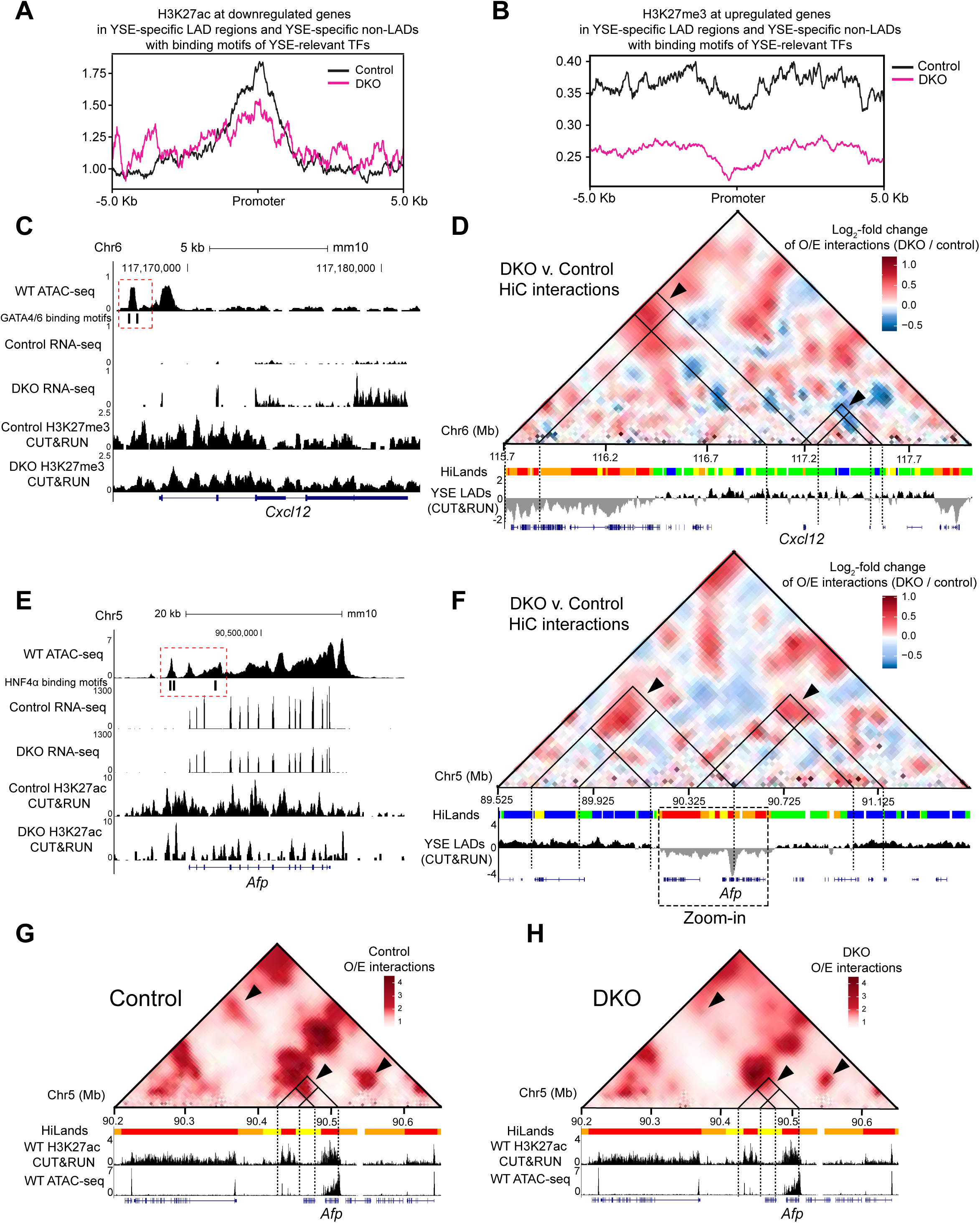
Lamin-A/B1 maintain 3D chromatin interactions to support gene expression regulated by YSE-relevant transcription factors. (A and B) Metaplots of H3K27ac CUT&RUN signal around promoters of downregulated genes associated with binding motifs of YSE-relevant transcription factors (A) and H3K27me3 CUT&RUN signal around promoters of upregulated genes associated with binding motifs of YSE-relevant transcriptions (B) in the indicated LAD and non-LAD regions. (C) Genome browser view of ATAC-seq peaks and GATA4/6 binding motifs around the upregulated broad-lineage gene *Cxcl12*, alongside tracks of bulk RNA-seq and H3K27me3 CUT&RUN from control and lamin-A/B1 DKO cells. An ATAC-seq peak containing GATA4/6 motifs is highlighted (dashed box). (D) Heatmap showing 3D chromatin interaction changes around *Cxcl12* upon lamin-A/B1 loss. Boxed regions and arrows highlight increased interactions with transcriptionally active HiLands-R and reduced interactions with repressive HiLands-B. (E) Genome browser view of ATAC-seq and HNF4α binding motifs around the downregulated gene *Afp* with known functions in YSE cells, alongside tracks of bulk RNA-seq and H3K27ac CUT&RUN from control and lamin-A/B1 DKO cells. ATAC-seq peaks containing HNF4α motifs are highlighted (dashed box). (F) Heatmap showing 3D chromatin interaction changes around *Afp* upon lamin-A/B1 loss. Boxed regions and arrows indicate increased interactions with repressive HiLands-B. (G and H) Heatmaps of observed/expected (O/E) Hi-C interactions of the zoomed-in region around *Afp* outlined by the dashed box in (F) of control (G) and lamin-A/B1 DKO (H) YSE cells. The boxed region and arrows highlight reduced interactions between *Afp* and transcriptionally active HiLands-R upon lamin-A/B1 loss.

To illustrate how lamin-A/B1 could coordinate with YSE-relevant transcription factors to repress gene expression, we examined the gene *Cxcl12*, encoding a chemokine known to regulate migration of many cell types^70^. *Cxcl12* expression is known to be promoted by GATA4 during various tissue development^71,72^. *Cxcl12* is upregulated upon lamin-A/B1 loss in YSE cells, and its promoter-proximal ATAC-seq peak contains binding motifs for YSE-relevant transcription factors GATA4 and GATA6 (Figure 6C, see dashed box). Furthermore, the *Cxcl12* gene is in a YSE-specific LAD region (Figure S6C) and is covered by repressive H3K27me3 modifications (Figure 6C), indicating its repression in YSE cells. Upon lamin-A/B1 loss, *Cxcl12* gains interactions with active HiLands-R regions and loses interactions with repressive HiLands-B (Figure 6D). Our H3K27me3 mapping showed that lamin-A/B1 deletion resulted in loss of the H3K27me3 modification around the *Cxcl12* promoter (Figure 6C). Together, these changes can result in increased GATA4 activity, thereby contributing to upregulation of *Cxcl12* upon lamin-A/B1 deletion.

Finally, we used the downregulated gene *Afp* to illustrate how lamin-A/B1 can coordinate with YSE-relevant transcription factors to support functional YSE gene expression. *Afp* is the major serum protein produced by YSE cells involved in transporting fatty acids and other nutrients to the embryo^73^. *Afp* expression in YSE cells is supported by the YSE-relevant transcription factor HNF4α^74^ and the locus is in a YSE-specific non-LAD flanked by shared LADs characterized by HiLands-B, -G, and -Y (Figure 3H, Figure S6D). Upon lamin-A/B1 deletion, the flanking LADs are detached as judged by their decreased C-scores (Figure S6D) and *Afp* gains interactions with the flanking repressive HiLands-B (Figure 6F). To facilitate visualization of chromatin interaction changes between *Afp* and nearby HiLands-R, we zoomed-in and visualized chromatin interactions immediately surrounding the *Afp* locus and nearby HiLands-R regions (see dashed box in Figure 6F enlarged in Figure 6G, 6H). This revealed a reduction of interactions between *Afp* and nearby accessible HiLands-R regions (Figure 6G, 6H). Consistent with these interaction changes, we found a loss of active enhancer regions at the *Afp* promoter and gene body as judged by a reduction of H3K27ac (Figure 6E). These changes could reduce HNF4α activity, thereby contributing to *Afp* downregulation.

## Discussion

Based on our findings, we propose a model where lamins maintain LAD organization and 3D chromatin interactions, which in turn influences lineage-relevant transcription factor function and gene expression programs (Figure 7). Our studies establish a conceptual framework for understanding how ubiquitous lamins regulate distinct transcriptional programs across diverse cell types during development.

**Figure 7.**
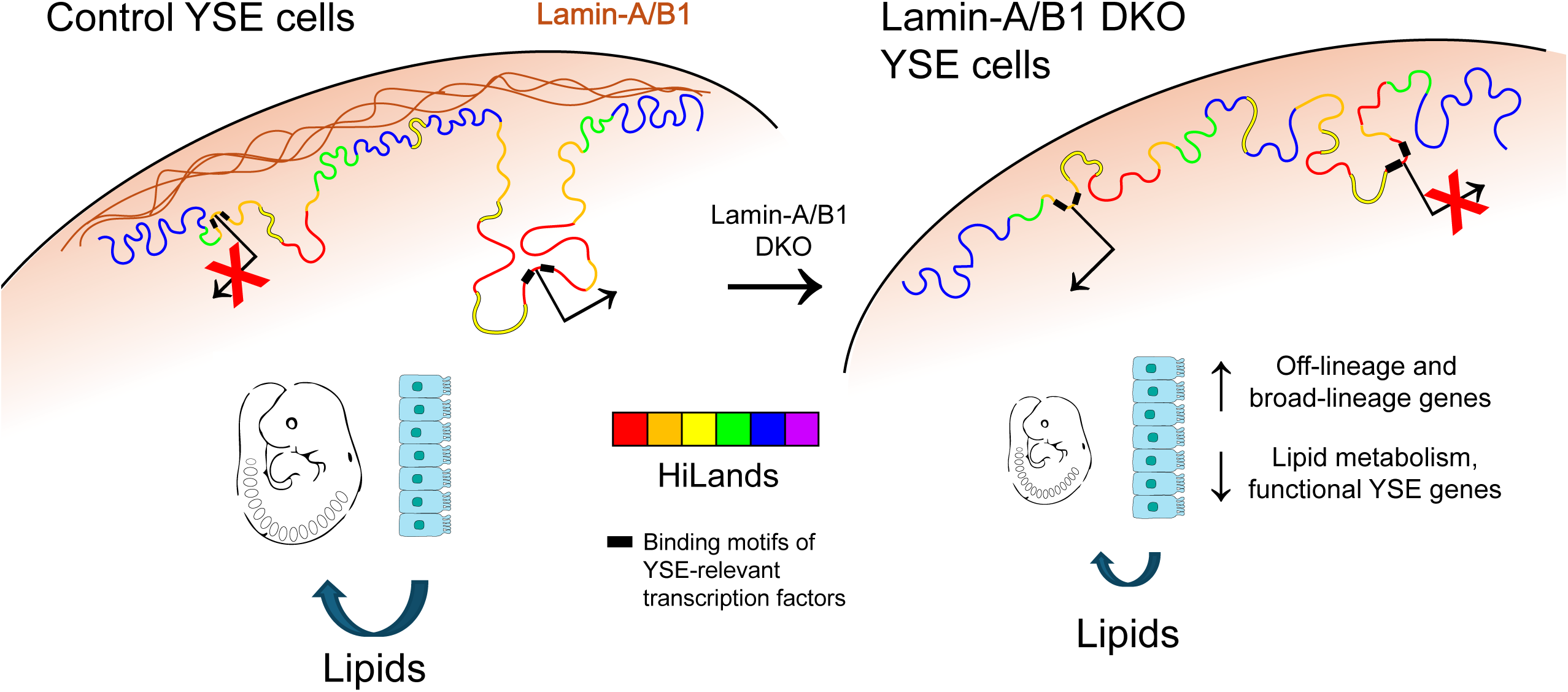
Model. Lamin-A/B1 maintain LADs characterized by HiLands-B, HiLands-G, and HiLands-Y in YSE cells, which orchestrates chromatin interactions and influences gene expression programs regulated by YSE-relevant transcription factors (binding motifs shown as black rectangles). By supporting YSE gene expression in collaboration with YSE-relevant transcription factors, lamin-A/B1 promote lipid transport to embryos while repressing off-/broad-lineage gene expression, thereby contributing to proper embryo growth and viability.

### The role of lamins during embryogenesis

Although deleting either A- or B-type lamins in mice results in defects of various organs, the mice survive until birth^28–32^. This has led to the prevailing view that lamins are not essential for embryogenesis and organogenesis. We show here that deleting lamin-A and -B1 results in an embryonic growth defect and midgestational lethality, demonstrating lamins are required for embryogenesis (Figure 1, S1). During midgestation, strong lamin-A expression is restricted to the extraembryonic organs, the yolk sac and placenta^17,18^, which support embryo growth. Since the embryonic growth defect upon lamin-A/B1 loss begins at E8.5, before the placenta becomes fully functional^42^, our findings suggest that these two lamins play a role in proper organogenesis and function of the yolk sac to support midgestational embryo development. Due to the difficulty of finding efficient Cre drivers to delete lamins in specific lineages at early developmental time points, we cannot exclude the possibility that lamin-A/B1 knockout also results in placental defects that may further impair embryo development. Therefore, to fully understand the role of lamins during embryogenesis, it is important to create effective Cre-drivers that allow timely knockout of different combinations of lamins in specific cell lineages.

It is challenging to study the developmental function of lamins because they are involved in both helping the nucleus withstand significant mechanical strain^31,32,34–36^ and supporting transcription via organizing LADs^21,23,24^. For cell lineages that are under large mechanical strain during embryogenesis, lamin knockout can cause DNA damage. For example, during embryonic brain development in mice, neural progenitor cells undergo interkinetic nuclear migration, which puts significant mechanical strain on the nucleus^75^. These cells require lamins to maintain their nuclear integrity, as either single- or double-knockout of lamin-B1 and -B2 results in DNA damage and apoptosis^31,32,35^. Since DNA damage leads to extensive downstream transcriptional changes and alters 3D genome organization^76,77^, it has been difficult to tease apart lamins’ direct role in maintaining 3D genome organization and transcription from the confounding effects of DNA damage.

The yolk sac is suspended in the amniotic fluid and is likely under low mechanical strain, which may explain why lamin-A/B1 deletion caused no increased DNA damage in YSE cells. The lack of DNA damage makes YSE cells a useful model to study lamins’ direct roles in genome organization and gene expression. Our study suggests it may be possible to find additional embryonic cell lineages under mild mechanical strain that do not suffer from DNA damage upon lamin loss. Further studying these cell types would enable investigation of lamins’ roles in genome organization and transcription during development of different lineages. Although lamins are ubiquitously expressed, their knockout leads to a wide range of phenotypes in different cell types^78–80^. Further understanding lamins’ role in maintaining the 3D genome organization of diverse lineages could shed light on how lamins maintain specific transcriptional programs which support diverse functions in different cell types.

### Developmental LAD remodeling and the role of lineage-relevant transcription factors

We show that YSE cells display more LAD formation than LAD detachment during development (Figure 3, S3), consistent with findings made in other differentiated cell types^14,57^. By coupling our LAD analyses with HiLands chromatin state modeling, we find the YSE-specific LAD regions and YSE-specific non-LADs contain genes with binding motifs of YSE-relevant transcription factors (Figure 3, S3). Our findings suggest that lineage-relevant transcription factors are involved in ensuring gene regulation in specific lineages by facilitating LAD detachment and preventing genes from becoming part of the lineage-specific LADs. Interestingly, in a melanotrope cell culture model, the melanotrope-associated transcription factor Pax7 was shown to mediate dissociation of enhancers from LAD-associated closed chromatin as mapped using lamin-B^81^. Furthermore, a recent preprint reported that limb-associated transcription factor motifs are enriched in regions that detach from the nuclear lamina during limb progenitor development^82^.

While lineage-relevant transcription factors may prevent LAD formation, the factors that promote lineage-specific LAD formation during development remain unclear. By annotating LADs with HiLands chromatin states, we were able to define regions within YSE-specific LAD regions that display increased lamin association. This enabled us to perform motif finding in these regions, and we found binding motifs of several transcription factors highly expressed in YSE cells with known repressive functions (Figure S3, Table S3). Interestingly, the transcriptional repressor cKrox has been shown to mediate LAD formation in tissue culture fibroblasts^83^. Therefore, transcriptional repressors expressed in a given cell lineage could play a role in establishing lineage-specific LADs during development. Future studies comparing LADs aided by HiLand modeling across many cell types could facilitate motif discovery and experimental validation of transcriptional repressors involved in lineage-specific LAD formation.

Although HiLands modeling has enabled us to identify lineage-relevant transcription factors that may aid lineage-specific LAD reorganization, how the constitutive LADs shared by most cell types are maintained is not well understood. Studies have shown that as LADs are mapped in increasing numbers of different cell types, the constitutive LAD regions shrink^13^. Thus, annotating LADs and modeling HiLands in a large number of cell types derived from different lineages may help to further narrow down the constitutive LADs regions and identify their DNA sequence features, which could aid the finding of factors that maintain these constitutive LADs.

### Lamins may collaborate with lineage-relevant transcription factors to achieve lineage appropriate gene expression programs during development

We find that lamin-A/B1 loss in YSE cells leads to downregulation of genes important for YSE function, alongside upregulation of off-lineage and broad-lineage genes (Figure 1, Table S1). Importantly, both the upregulated and downregulated genes in YSE cells are enriched for binding motifs of transcription factors involved in YSE lineage development, suggesting lamins may coordinate with lineage-relevant transcription factors to regulate gene expression. Our findings offer an explanation for how the ubiquitously expressed lamins may support cell-type-specific gene expression. An important question is whether the findings reported here only apply to YSE cells or reflect a general principle that enables lamins to support different gene expression programs in diverse cell lineages. To explore this, it will require one to use lineages not under intense mechanical strain that therefore do not suffer from DNA damage upon lamin loss. Additionally, while the yolk sac offers a rare opportunity for the isolation of pure YSE cells during embryogenesis with simple mechanical dissection, isolating sufficient numbers of pure cells from most tissues is challenging. It would be important to create lineage-specific fluorescent reporter mouse models to enable efficient isolation of a pure cell lineage for this kind of study.

### Lamins may influence lineage-relevant transcription factors by maintaining chromatin neighborhoods

By utilizing the HiLands framework, we find lamin-A/B1 maintain organization of LADs characterized by HiLands-B, HiLands-G, and HiLands-Y in YSE cells (Figure 4, S4). Disruption of these LAD regions correlates with chromatin neighborhood changes and dysregulation of genes containing YSE-relevant transcription factor motifs. Importantly, HiLands-based analyses revealed gene downregulation correlates with decreased interactions with active enhancers while gene upregulation correlates with decreased interactions with repressive LAD chromatin (Figure 5, S5, 6, S6). These findings provide a rationale for how lamins can influence lineage-relevant transcription factor function by maintaining proper chromatin neighborhoods to both support functional gene expression and repress off-lineage and broad-lineage genes in a given cell lineage. Testing the causal relationship among LAD organization, chromatin interactions, transcription factor function, and gene expression remains challenging due to the large genomic size of LADs and the extensive chromatin interaction changes that accompany lamin loss. Future studies to test causality may benefit from recently developed CRISPR-based genome tethering approaches^84^. Such methods could enable the repositioning of a given detached LAD back to the nuclear lamina, allowing direct testing of whether restoring a given LAD rescues chromatin interactions along with correcting transcription factor function for the corresponding gene(s). In addition, tethering of individual genes back to their proper active or repressive chromatin environments could help determine how local chromatin context influences transcription factor function and gene regulation.

By linking lamins’ role in 3D genome organization to transcriptional programs mediated by lineage-relevant transcription factors, our study offers a conceptual framework to further interrogate how lamins exert cell-type-specific effects in diverse lineages during development. Since lamins and nuclear lamina organization are frequently disrupted during aging and in disease^85^, it will be important to investigate how these changes could influence chromatin architecture and transcription factor-mediated gene regulation, potentially contributing to pathological gene expression programs.

### Limitations of the study

While the transcriptional defects in YSE cells upon lamin-A/B1 loss can help explain the embryonic growth defect and lethality, we did not directly test whether YSE dysfunction causes embryo lethality. Due to inefficiency of the *Ttr*-Cre driver, we were unable to achieve effective conditional knockout of lamin-A/B1 in YSE cells by E9.5. Since the placenta can take-over nutritive functions and mask embryonic phenotypes around this timepoint^42^, additional more efficient Cre-lines need to be developed to further confirm the YSE-specific function of lamin-A/B1. Although our genomics study is correlative, it offers a clear path forward to develop experimental systems to address causality. We note the chromatin tether systems we discussed will take a long time to implement and optimize in mice. In addition, while we used motif analysis as a proxy for transcription factor binding, it would be ideal to verify transcription factor binding by CUT&RUN. However, such experiments are currently challenging due to limited yields of YSE cells and labile association of transcription factors with chromatin, making CUT&RUN mapping difficult. Further methodology development, along with studies in other cell types yielding higher cell numbers, should help overcome these limitations.

## Acknowledgements

We thank Allison Pinder, Fred Tan, and Javier Carpinteyro Ponce for help with sequencing; Mahmud Siddiqi for microscopy and Imaris analysis assistance; Lynne Hugendubler for technical assistance; Joseph Tran, Katherine Bossone, and Ross Pedersen for technical advice and helpful feedback; and additional members of the Zheng lab for helpful discussion. This work was supported by R01GM106023 (Y. Zheng, R.D. Goldman), R01GM110151 (Y. Zheng), 1R01GM157598-01 (Y. Zheng), RS-2023-00222784 (Y. Kim), RS-2025-25441283 (Y. Kim) and RS-2024-00437643 (Y. Kim).

## Author Contributions

S.D. performed experiments, bioinformatic analysis, visualization, and writing of the original draft. X.Z. advised on bioinformatic analyses. J.H. performed and analyzed lamin TKO embryo dissections. Y.K., L.K., and R.M. developed low-cell Hi-C methodology. Y.Z. and S.D. performed writing and editing. Y.Z. and Y.K. supervised aspects of the work and acquired funding. Y.Z. and S.D. conceptualized the study.

## Declaration of AI

During preparation of the manuscript, S.D. utilized ChatGPT-5 mini (OpenAI) to assist with editing sections of the text for clarity. All content was reviewed and edited by the authors. Y.Z. and S.D. utilized Zo Computer (https://www.zo.computer) to aid literature survey and confirmation of certain past research findings.

## Methods

## RESOURCE AVAILABILITY

### Lead Contact

Further inquiries should be directed to the corresponding authors, Sara Debic and Yixian Zheng.

### Materials availability

Mouse lines used to generate the genotype combinations in this study are available from Jax, the International Mouse Strain Resource, the Mutant Mouse Resource and Research Centers, and Charles River (see Methods for strain details).

### Data and code availability

RNA-seq, ATAC-seq, CUT&RUN, and Hi-C datasets generated in this study have been deposited in GEO under SuperSeries accession number GSE318232 and accession numbers GSE318087, GSE318088, GSE318089, GSE318100, and GSE318101 included therein. Custom code used in this study will be made publicly available at https://github.com/saradebic/lamin-YSE-multiomics.

## KEY RESOURCES TABLE

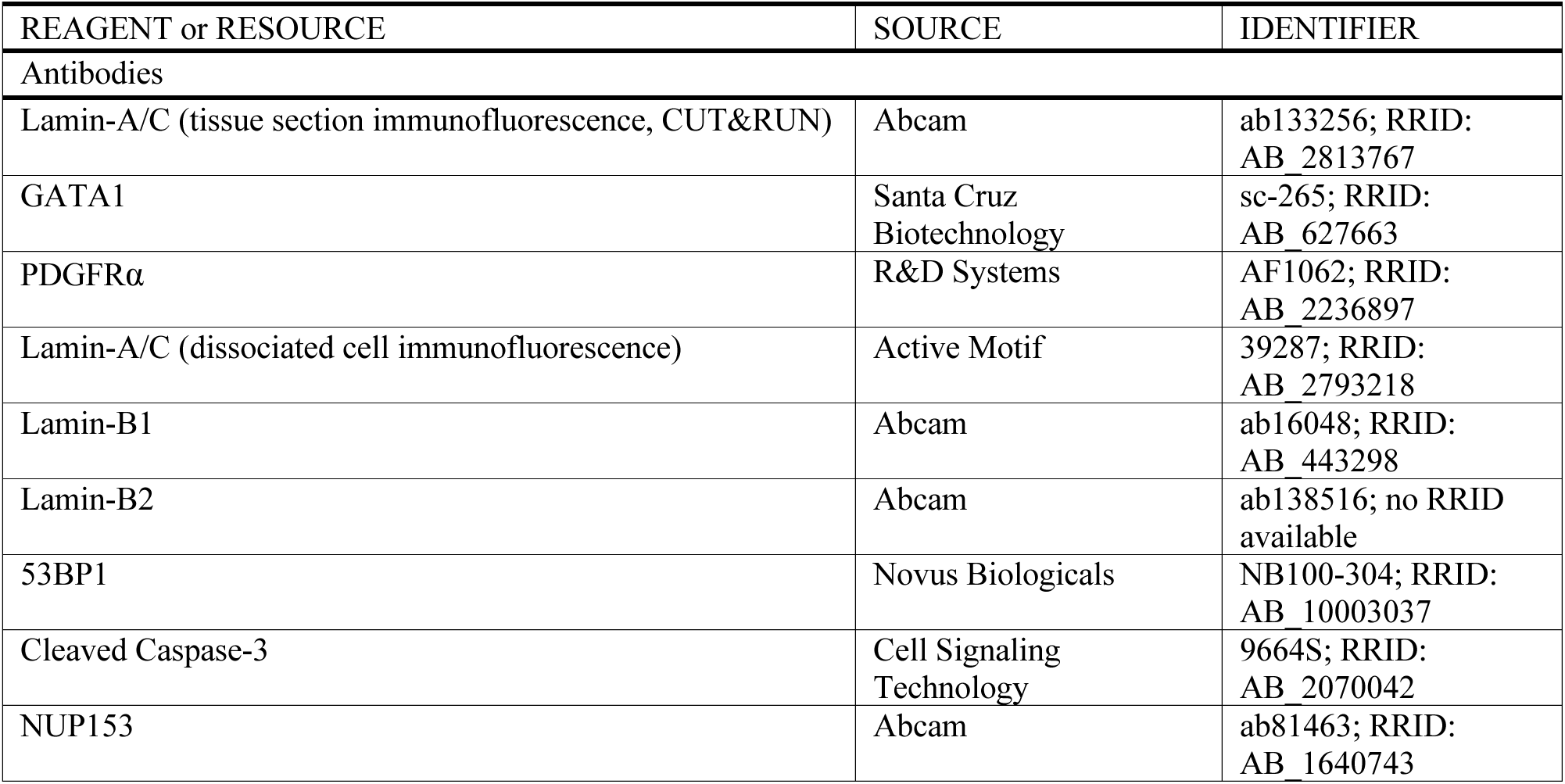
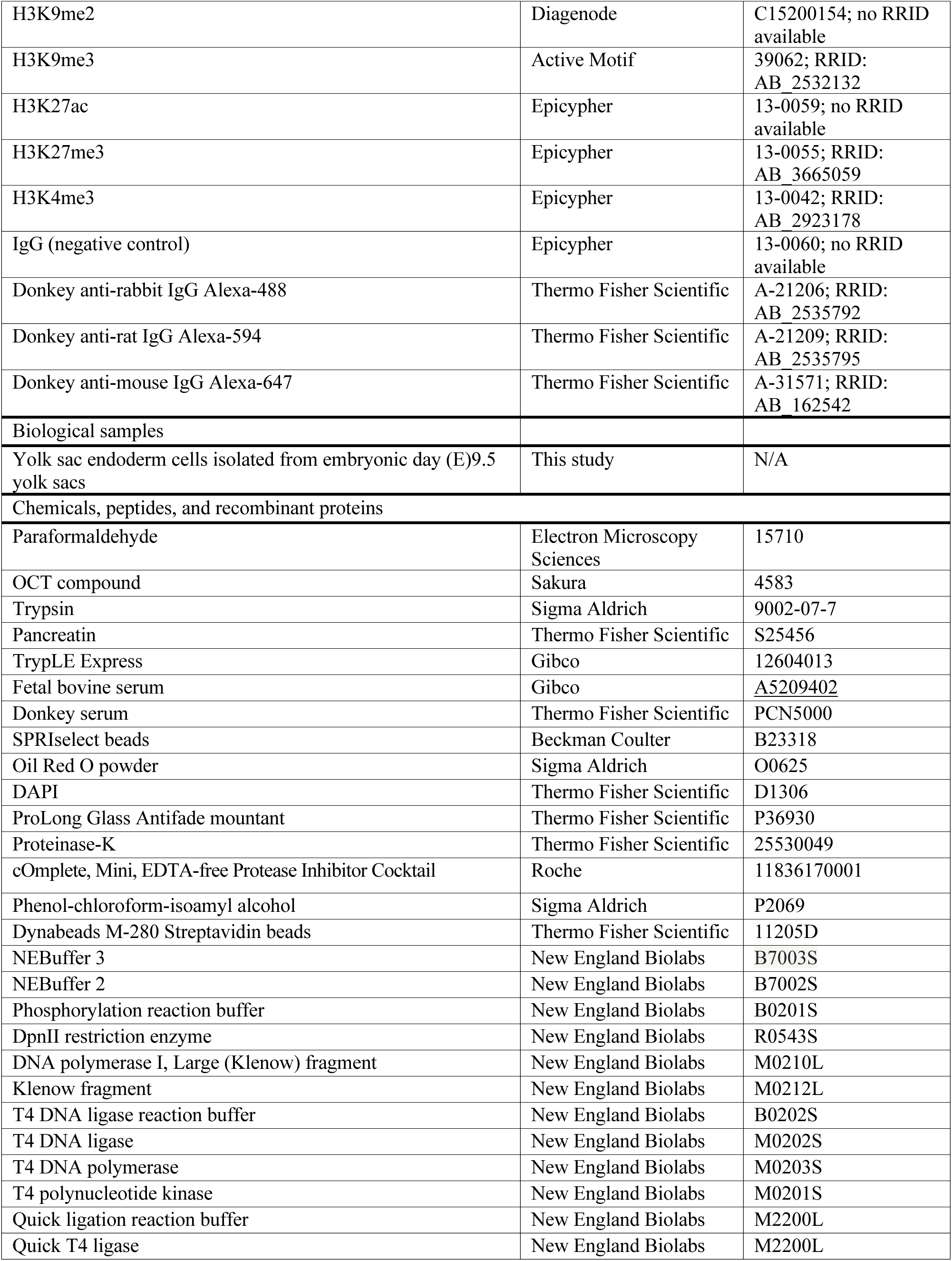
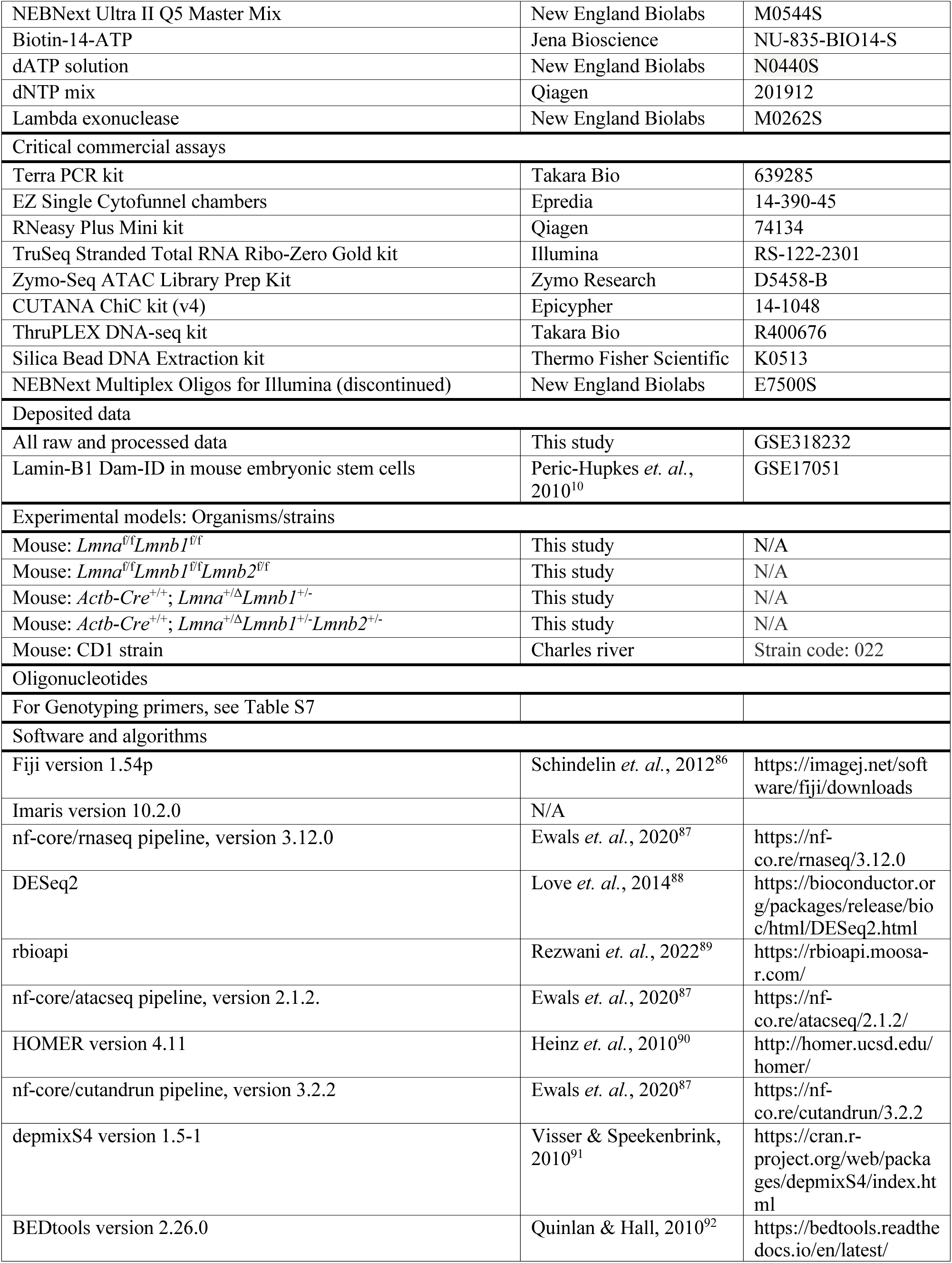
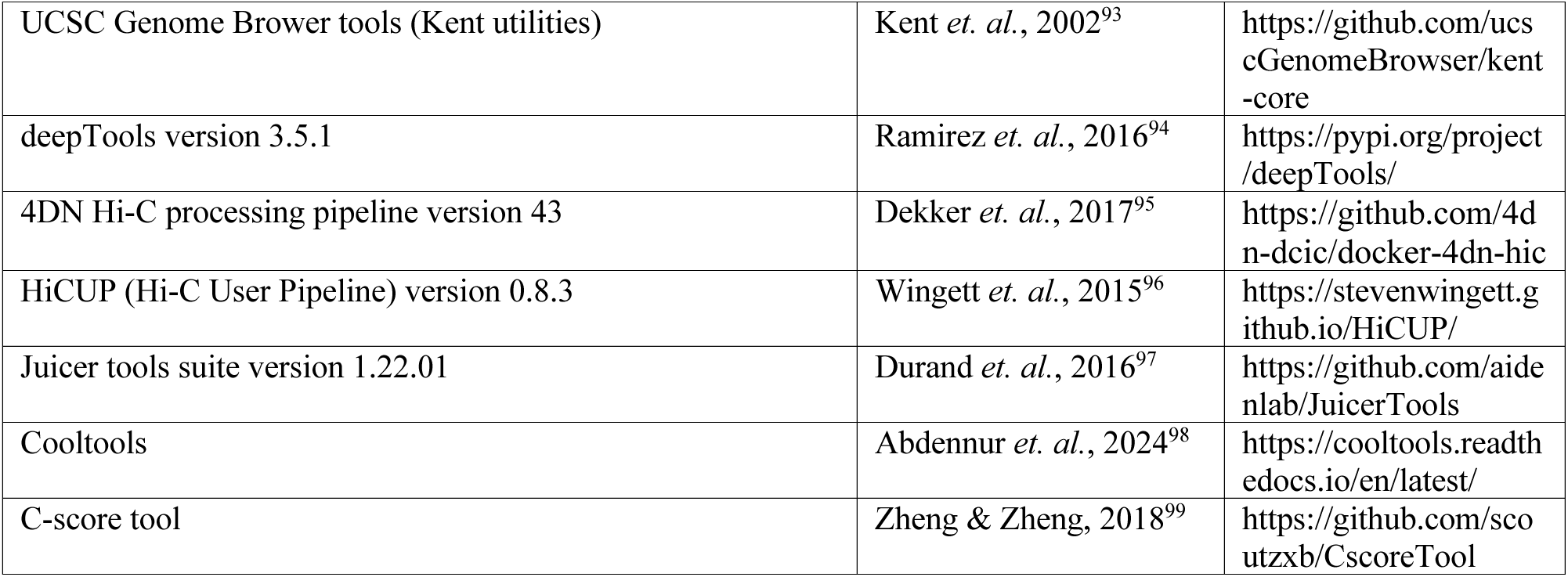

## EXPERIMENTAL MODEL AND SUBJECT DETAILS

### Mouse husbandry

All experimental mice were housed in a facility on a 12h light/dark cycle at 20-22°C with 30-50% humidity. Bedding, food, and water were routinely changed. Animal health was monitored using a sentinel surveillance program in which CD1 sentinel mice were exposed to soiled bedding from colony cages and tested quarterly for viral and parasitic pathogens by serology and PCR at Charles River Laboratories. All animal procedures were approved by the Institutional Animal Care and Use Committee (IACUC) and were conducted in accordance with NIH and institutional guidelines.

### Mouse strains

To generate lamin-A/B1 double-knockout (DKO) and lamin triple-knockout mouse embryos, conditional alleles for *Lmna*, *Lmnb1*, and *Lmnb2* were used. A *Lmna*^f/f^*Lmnb1^f/f^* line was generated by crossing *Lmnb1^f/f^*mice carrying the *Lmnb1*^tm1a(EUCOMM)Wtsi^ allele (EUCOMM project, International Mouse Strain Resource) with *Lmna^f^*^/f^ mice (JAX stock #026284). For the lamin triple-flox line, the *Lmnb2^f/f^* allele was derived from *Lmnb2*^tm1a(KOMP)Wtsi^ (KOMP project, International Mouse Strain Resource) and the neomycin cassette flanked by FRT sites was removed by breeding with *Actb*-FLPe mice. *Actb-Cre^+/+^*; *Lmna^+^*^/Δ^*Lmnb1^+^*^/-^ and *Actb-Cre^+/+^*; *Lmna^+/^*^Δ^*Lmnb1^+^*^/-^*Lmnb2^+/-^*lines were generated by crossing mice heterozygous for *Lmna*, *Lmnb1,* and *Lmnb2* alleles generated previously^30,31^ (*Lmnb1* and *Lmnb2* deletion alleles available at Mutant Mouse Resource and Research Centers ID #42096) with mice homozygous for Cre recombinase driven by the human β-actin promoter (JAX stock #019099). Both lines were bred to homozygosity for the *Actb-Cre* allele. *Lmna*^Δ/Δ^*Lmnb1^-^*^/-^ embryos were obtained by crossing *Lmna^f^*^/f^*Lmnb1^f^*^/f^ females with *Actb-Cre^+/+^*; *Lmna^+^*^/Δ^*Lmnb1^+^*^/-^ males. Similarly, *Lmna*^Δ/Δ^*Lmnb1^-^*^/-^*Lmnb2^-/-^*embryos were obtained by crossing *Lmna^f^*^/f^*Lmnb1^f^*^/f^*Lmnb2^f/f^*females with *Actb-Cre^+/+^*; *Lmna^+/^*^Δ^*Lmnb1^+^*^/-^*Lmnb2^+/-^* males. All conditional knockout and Cre-driver lines were maintained in a mixed genetic background of CD1, 129Sv, and C57BL/6J strains. Embryos of both sexes were used, and sex was not considered as a biological variable in this study. For all experiments involving *Lmna*^Δ/Δ^*Lmnb1^-^*^/-^ yolk sac endoderm (YSE) cells, *Lmna^+/^*^Δ^*Lmnb1^+^*^/-^ YSE cells were used as controls. For all experiments using wild-type YSE cells, YSE cells were isolated from yolk sacs of CD1 mouse embryos.

## METHOD DETAILS

### Embryo collection and genotyping

Pregnant females were euthanized at embryonic day 9.5 (E9.5), E8.5, or E7.5, with the day of vaginal plug detection designated as E0.5. Decidua were isolated from the uteri of pregnant females, and embryos with intact yolk sacs were dissected out of decidua in phosphate-buffered saline (PBS) using Dumont #5 fine-tipped forceps under a Leica M125 dissecting microscope. Embryos were staged according to Theiler staging criteria based on limb morphology and somite number and were imaged using a Leica M125 dissecting microscope equipped with a Leica IC80 HD camera. Whole embryos were used for genotyping using PCR. Briefly, whole embryos were lysed in 300 μL of 50 mM sodium hydroxide and incubated at 90°C for 10 minutes. Embryo lysates were neutralized with 30 μL of Tris-HCl (pH = 8.0) and centrifuged at maximum speed for 1 minute in a tabletop centrifuge. 1 μL of supernatant was used as template DNA for PCR using the Terra PCR kit following the manufacturer’s protocol. PCR cycling conditions were as follows: 98°C for 2 minutes, followed by 35 cycles of 98°C for 10 seconds, 63°C for 15 seconds, and 68°C for 1 minute. All locus-specific genotyping primers are listed in Supplementary Table S7, and PCR products were visualized on 2% agarose gels.

### Yolk sac endoderm cell isolation and dissociation

We adapted a published protocol^45^. Briefly, to isolate yolk sac endoderm tissue at E9.5, yolk sacs were separated from embryos using Dumont #5 fine-tipped forceps and incubated in an enzyme solution containing 5 mg/mL trypsin and 25 mg/mL pancreatin in PBS for 4 hours at 4°C with gentle shaking. Following enzymatic treatment, the visceral yolk sac endoderm layer was manually separated from the yolk sac (extraembryonic) mesoderm layer using Dumont #5 fine-tipped forceps under a Leica M125 dissecting microscope. To obtain a single-cell dissociation, isolated yolk sac endoderm cell tissue was incubated in 1X TrypLE Express (Gibco) at 37°C for 7 minutes followed by gentle mechanical trituration using a P200 micropipette. This enzymatic and mechanical dissociation cycle was repeated two to three times, or until the tissue appeared fully dissociated into single cells. Enzymatic activity was quenched by the addition of fetal bovine serum (FBS; Gibco) to a final concentration of 10% (volume/volume).

### Cryosectioning

Intact decidua containing embryos and yolk sacs were isolated at E9.5 and fixed in 4% paraformaldehyde (PFA) in PBS overnight at 4°C, followed by three washes in PBS. Next, the decidua were incubated in 30% sucrose solution in PBS overnight, and then embedded in OCT compound (Sakura, 4583) in cryomolds, and frozen at -80°C. Cryosectioning was performed using a Leica CM3050 S cryostat at −20°C. Decidua were sectioned into tissue slices ranging from 7 to 15 μm in thickness and mounted onto Superfrost Plus glass slides (VWR). These tissue slices included cross sections of the yolk sac and embryo.

### Genotyping from cryosections

Genotyping was performed directly from tissue sections by scraping embryonic material into 0.1% Tween-20 in PBS, followed by lysis in RIPA buffer (50 mM Tris-HCl, 150 mM NaCl, 0.1% SDS, 0.5% sodium deoxycholate, 1% Triton-X-100, pH = 7.5) containing 40 μg of proteinase K. Samples were incubated overnight at 50°C with shaking. Genomic DNA was purified the following day using SPRIselect beads (Beckman Coulter) following the manufacturer’s protocol, and 1 μL of purified DNA was used as template for genotyping PCR (described above).

### Lamin immunofluorescence of yolk sac tissue sections

Yolk sac tissue sections were fixed using paraformaldehyde as described above, and sections were outlined with a PAP pen to restrict staining solutions. To enhance antibody penetration into yolk sac tissue, the tissue sections were treated with TrypLE solution for 10 minutes at room temperature prior to permeabilization. Samples were permeabilized with 0.25% Triton-X-100 for 20 minutes at room temperature and subsequently incubated overnight at 4°C in a humidified chamber with primary antibody diluted in block solution (10% donkey serum volume/volume, 10% BSA weight/volume, 10 mM sodium azide, and 0.1% Tween-20 in 1X PBS volume/volume). Immunofluorescence staining was performed using primary antibodies against lamin-A/C (Abcam, ab133256), GATA1 (Santa Cruz Biotechnology, sc-265), and PDGFRα (R&D systems, #AF1062) each diluted 1:100 in block solution. GATA1 and PDGFRα staining was included to identify red blood cells and YSE cells, respectively. The following day, samples were washed three times with PBS (5 minutes each) and incubated with a 1:1000 dilution of Alexa Fluor-conjugated secondary antibodies (ThermoFisher) in block solution containing 5 μg/uL DAPI (ThermoFisher, D1306) for one hour at room temperature protected from light. After an additional PBS wash, the sections were mounted with ProLong Glass Antifade Mountant (ThermoFisher, P36930) and imaged using a Leica SP5 confocal microscope.

### Lamin immunofluorescence of cytospun YSE cells

YSE cells were isolated from lamin-A/B1 DKO and control littermates as described above. Briefly, YSE cells were dissociated in 125 μL of TrypLE Express, quenched with an equal volume of DMEM cell culture media, and fixed in 2% paraformaldehyde in PBS for 10 minutes at room temperature. The fixation was quenched by adding glycine to a final concentration of 125 mM. To spin the YSE cells down onto glass slides, the fixed cells were loaded into EZ Single Cytofunnel chambers (Epredia, 14-390-45) and centrifuged onto Superfrost Plus glass slides (VWR) in a Cytospin 4 centrifuge (Thermo Scientific, TH-CYTO4) for 10 minutes at 300 x g. Immunofluorescence staining for lamin-A/C, lamin-B1, and lamin-B2 was performed as described above. The following primary antibodies were used: lamin-A/C (Active Motif, 39287; 1:200 dilution), lamin-B1 (Abcam, ab16048; 1:200 dilution), and lamin-B2 (Abcam, ab138516; 1:200 dilution).

### Immunofluorescence for apoptosis, DNA damage, and nuclear pore complex markers

Yolk sac tissue sections from lamin-A/B1 DKO and littermate controls were used to assess levels of DNA damage, apoptosis, and nuclear pore complex clustering in YSE cells upon lamin-A/B1 loss. DNA damage was assessed by performing immunofluorescence for 53BP1 (Novus, NB100-304; 1:1000 dilution), apoptosis was assessed by immunofluorescence against cleaved caspase-3 (Cell Signaling, #9664S; 1:200 dilution), and nuclear pore complexes were visualized by staining for NUP153 (Abcam, ab81463; 1:200 dilution). Immunofluorescence of yolk sac tissue sections was performed as described above. YSE cells were identified based on their characteristic morphology within the yolk sac, including nuclear size, shape, and position. To quantify 53BP1 and cleaved caspase-3 foci, we used the Analyze Particles function in Fiji (ImageJ) after performing manual thresholding. 53BP1 foci were quantified per DAPI-stained YSE cell nucleus, and cleaved caspase-3 foci were assigned to the nearest DAPI-stained YSE cell nucleus based on visual inspection of merged channels. Particles with a size between 0.1 and 100 μm^2^ and with a circularity between 0.3 and 1 were included in both analyses, with identical thresholding and particle size parameters applied across all images.

### Oil Red O lipid staining

Embryo cryosections from lamin-A/B1 DKO and littermate control embryos were used for Oil Red O staining. A stock Oil Red O solution was prepared by dissolving 300 mg of Oil Red O powder (Sigma) in 100 mL of isopropanol. The working Oil Red O solution was made by mixing 30 mL of stock solution with 20 mL of distilled water and filtering through a 0.45 um filter before use. Air dried tissue sections were first equilibrated in 60% isopropanol diluted in distilled water for 1 minute using a coplin jar. Next, the sections were stained in a working Oil Red O solution for 15 minutes using a coplin jar. Lastly, the slides were washed in 60% isopropanol once again, and then in distilled water briefly. The slides were mounted with ProLong Glass Antifade Mountant (ThermoFisher, P36930) and imaged on a Nikon Ti2 inverted microscope equipped with a Hamamatsu ORCA-Flash4.0 V3 camera using a 10X objective. The number of Oil Red O particles in lamin-A/B1 DKO and control embryo sections was quantified using the Analyze Particles function in Fiji (ImageJ). Images were converted to 8-bit and inverted prior to manual thresholding. Particles with a size between 0.1 and 10 μm^2^ and with a circularity between 0.3 and 1 were included in this analysis, with identical thresholding and particle size parameters applied across all images.

### Bulk RNA-sequencing (RNA-seq)

Total RNA was isolated from lamin-A/B1 DKO and control YSE cells using the RNeasy Plus Mini kit (Qiagen, 74134) following the manufacturer’s protocol. Briefly, intact yolk sac endoderm cell layers (not dissociated) were dissected as outlined above, transferred to microcentrifuge tubes, and flash-frozen for 5 seconds in liquid nitrogen. Each sample was then lysed in 350 μL of Buffer RLT Plus and homogenized by passing the lysate through a 20-gauge needle attached to a sterile syringe at least three times. RNA was purified per the kit protocol and eluted in 20 μL of RNase-free water. Ribosomal RNA was depleted using the TruSeq Stranded Total RNA Ribo-Zero Gold kit (Illumina, RS-122-2301). Libraries were sequenced on an Illumina NextSeq 500 platform using 50 base pair (bp) single-end reads.

### ATAC-seq

ATAC-seq was performed using the Zymo-Seq ATAC Library Prep Kit (Zymo, D5458-B) according to the manufacturer’s protocol. 50,000 live YSE cells were isolated from a single E9.5 CD1 yolk sac and dissociated as described above. Cell viability and counts were assessed using Trypan Blue staining and a hemocytometer. Nuclei were isolated following the Zymo-Seq protocol and incubated with the supplied Transposition Mix (containing Tn5 transposase and buffer) at 37°C for 30 minutes. Following transposition, DNA was purified using Zymo-Spin IC Columns included in the kit. Sequencing libraries were constructed using the kit’s library prep reagents and DNA was amplified with 10 cycles of PCR as recommended by the manufacturer. Final libraries were sequenced on the Illumina NextSeq 500 with paired-end 50/50 bp read length.

### CUT&RUN for mapping LADs and histone post-translational modifications

CUT&RUN was performed using the CUTANA ChiC kit (Epicypher, Version 4, Cat. No. 14-1048) following the manufacturer’s protocol. YSE cells isolated from one E9.5 CD1 strain yolk sac were used for each CUT&RUN targeting individual lamin proteins (lamin-A, lamin-B1, lamin-B2) or individual histone modifications (H3K4me3, H3K27ac, H3K27me3, H3K9me2, H3K9me3). CUT&RUN for H3K27me3 and H3K27ac was also performed on lamin-A/B1 DKO and control YSE cells. Briefly, dissociated YSE cells were washed once with Wash buffer, then incubated with activated Concanavalin A (ConA) beads for 10 minutes. After adsorption of cells to magnetized ConA beads, the supernatant was removed using a magnetic stand and 50 μL of Antibody buffer was added to the cells. For H3K4me3 and control IgG CUT&RUN, antibodies supplied in the CUTANA kit were used. The following antibodies were used for additional CUT&RUN: Lamin-A/C (Abcam, ab133256), Lamin-B1 (Abcam, ab16048), Lamin-B2 (Abcam, ab138516), H3K9me2 (Diagenode, C15200154), H3K9me3 (Active Motif, Cat. no. 39062), H3K27ac (Epicypher, 13–0059), H3K27me3 (Epicypher, 13-0055). For each sample, 1 μL of the respective antibody was used. For validation of histone modification antibodies, 2 μL of the K-MetStat Panel (Epicypher) was added at a dilution of 1:10. Cells were incubated with antibody overnight at 4°C on a nutator set to rotate no further than a 60-degree angle. The following day, cells were washed twice with 200 μL of Permeabilization buffer, then resuspended in 50 μL of Permeabilization buffer containing 2.5 μL of pAG-MNase. After a 10-minute incubation at room temperature, cells were again washed twice with 200 μL of Permeabilization buffer and resuspended in 50 μL of Permeabilization buffer with 1 μL of 100 mM calcium chloride added to initiate digestion. The reactions were incubated at 4°C for two hours on a nutator with the same settings as above. To stop the reaction, Stop Master Mix containing 1 μL of 1:25 diluted *E. coli* Spike-in DNA from the kit was added. The reactions were incubated at 37°C for 10 minutes in a thermocycler, and afterwards the supernatants were collected for DNA isolation. DNA was purified using 119 μL of SPRIselect beads included in the kit following standard bead cleanup protocols. Sequencing libraries were prepared using the Takara ThruPLEX DNA-seq kit according to the manufacturer’s protocol using ∼1 ng of CUT&RUN DNA as input with 12 cycles of PCR amplification. Final libraries were sequenced with the Illumina NextSeq 500 with paired-end 50/50 bp read length.

### Hi-C

#### 1. YSE cell harvest and crosslinking

Lamin-A/B1 DKO and control YSE tissues were isolated and dissociated into single cells as outlined above. YSE cells from one yolk were used per biological replicate. Hi-C was performed using an optimized low-input protocol with carriers; detailed methodological description will be reported elsewhere and will be provided upon request. Briefly, YSE cells were fixed with 2% formaldehyde for 10 minutes at room temperature. Formaldehyde was quenched by the addition of glycine to a final concentration of 125 mM, followed by incubation for 5 minutes at room temperature and 15 minutes on ice. Cells were then pelleted by centrifugation, washed with 1 mL of ice-cold PBS, and centrifuged again under the same conditions.

#### 2. Cell lysis and enzyme digestion

The YSE cells were resuspended in 1 mL of lysis buffer consisting of 10 mM Tris-HCl at pH=8, 10 mM NaCl, 0.2% IGEPAL CA-639 (NP-40), and 1X protease inhibitor cocktail (Roche, 11836170001) and rotated for 30 minutes at 4°C. After lysis, cells were centrifuged for 8 minutes at 1,000 x g at 4°C and washed with 500 μL of cold 1.2x NEB3 buffer (NEB, B7003S). After another centrifugation at the same conditions, the pellet was resuspended in 40 μL NEB3 buffer, and 1.2 μL of 10% SDS was added (final concentration of SDS 0.3%). The reaction was incubated at 65°C for 10 minutes using a PCR machine, and afterwards immediately put on ice. The SDS was quenched by the addition of 4 μL of 20% Triton X-100 and carefully mixing the reaction by inverting the tube. The reaction was incubated at 37°C for one hour, followed by the addition of 2.5 μL of DpnII restriction enzyme (NEB, R0543S). The first restriction enzyme incubation was 90 minutes at 37°C, with gentle mixing by tapping the tube every half hour. Following this, another 2.5 μL of DpnII was added and incubated at 37°C for 90 minutes with gentle mixing every half hour.

#### 3. Biotin filling and proximity ligation

To fill-in the digested DNA ends with biotin, we prepared a reaction mixture by adding 0.5 μL of a 10 mM dCTP, dGTP, and dTTP mix (Qiagen, 201912), 1 μL of 1 mM biotin-14-ATP (JenaBioSciene, NU-835-BIO14-S), and 1.5 μL of Klenow fragment (NEB, M0210L 5 units/μL) to the DpnII reaction. The biotinylation reaction was incubated at 37°C for 60 minutes and afterwards put on ice. The blunted DNA was ligated by the addition of 50 μL of 10x T4 ligase buffer (NEB, B0202S), 5 μL of 100x BSA (NEB, 20 mg/mL), 390.8 μL of water, and 1 μL of NEB T4 ligase (NEB, M0202S, 400 units/μL). The ligation reaction was incubated overnight at 16°C without shaking.

#### 4. Decrosslinking and DNA purification

To reverse crosslinking, 2.5 μL of 20 mg/mL proteinase K (ThermoFisher, 25530049) was added and incubated overnight at 65°C. The next day, an additional 2.5 μL of 20 mg/mL proteinase K was added and incubated at 65°C for two hours. For DNA extraction, we added 500 μL of phenol-chloroform-isoamyl alcohol at a 25:24:1 ratio (Sigma, P2069) and vortexed vigorously. After centrifuging for 5 minutes at 16,000 x g at room temperature, we removed the supernatant to a new Eppendorf tube and repeated the phenol-chloroform-isoamyl DNA extraction process one more time. For additional DNA purification, we used the Silica Bead DNA Extraction kit (ThermoFisher, k0513) according to the manufacturer’s instructions. Briefly, we mixed 500 μL of phenol-chloroform-isoamyl alcohol extracted DNA with 1.5 mL of silica bead binding buffer and added 10 μL of silica bead suspension. The mixture was incubated at 55°C for 15 minutes, with occasional mixing. The silica beads were spun down by centrifuging briefly in a tabletop centrifuge at high speed for 5 seconds, after which the supernatant was carefully removed. The beads were then washed three times with 500 μL of ice-cold wash buffer included in the kit. The silica bead pellet was then air dried completely for at least 30 minutes after which it was resuspended in 45 μL of elution buffer (10 mM Tris, pH = 8) and incubated at 55°C for 5 minutes. Finally, the silica beads were spun down as outlined above and the supernatant was transferred into a Spin-X tube filter (Sigma, CLS8162 or Costar, 8160) to remove the residual silica beads. The solution in Spin-X tubes was centrifuged at 16,000 g for 5 minutes at 4°C and 40 μL of the filtered DNA solution was used for the de-biotinylation reaction.

#### 5. Removing biotin from un-ligated DNA

The filtered DNA solution was mixed with 1 μL of 10 mg/mL of bovine serum albumin, 10 μL of 10x NEB2 buffer (B7002S), 12.5 μL of 10 mM dNTP mix, 1.6 μL of T4 DNA polymerase (M0203S, 3 U/μL), and 34.9 μL of distilled water. The reaction was incubated for 2 hours at 20°C without shaking, and the reaction was stopped by adding 2 μL of 0.5 M EDTA. The DNA was purified from the reaction using the silica bead DNA extraction kit, as outlined above except the DNA was eluted in 50 μL of elution buffer. 50 μL of DNA was used for sonication.

#### 6. Sonication and DNA end repair

50 μL of the DNA suspension was transferred to a microTUBE AFA Fiber Screw-cap (Covaris, 520166) for sonication with the Covaris M220 instrument. Hi-C DNA was sheared for 90 seconds at 20°C with a peak incident power of 50 Watts, duty cycle of 20%, and 200 cycles per burst. End repair of the sonicated DNA was performed by mixing 50 μL of sonicated Hi-C DNA with 10 μL of 10x phosphorylation reaction buffer (NEB, B0201S), 0.5 μL of T4 DNA polymerase (NEB, M0203S), 0.5 μL of T4 polynucleotide kinase (NEB, M0201S), 5 μL of 10 mM dNTP mix (Qiagen, 201912), 15 μL of 10 mM dATP (NEB, N0440S), 2 μL of 10x diluted Klenow fragment DNA polymerase I (NEB, M0210L), and 17 μL of distilled water. The DNA end repair reaction was incubated at 37°C for 30 minutes in a PCR machine.

#### 7. Streptavidin capture

For all streptavidin capture steps, we used low-adhesion Eppendorf tubes. 20 μL of resuspended Dynabeads M-280 streptavidin beads (ThermoFisher, 11205D) were aliquoted in an Eppendorf tube. The beads were washed with 100 μL of 2x BW buffer consisting of 10 mM Tris-HCl with a pH of 7.5, 1 mM EDTA, and 2 mM NaCl. A magnetic stand was used to remove the supernatant from magnetic streptavidin beads at all wash steps. 100 μL of the end-repaired Hi-C DNA was mixed with 100 μL of 2x BW and added to the pre-washed streptavidin beads. The beads were resuspended by mild pipetting and rotated for 60 minutes at room temperature. The supernatant was subsequently removed with the aid of a magnetic stand and the beads were washed twice with 250 μL of 1x BWT buffer consisting of 5 mM Tris-HCl at pH 7.5, 0.5 mM EDTA, 1M NaCl, and 0.1% of Triton-X-100. Then, the beads were transferred to new tubes and washed twice with 250 μL of 10 mM Tris-HCl at a pH of 8.0. The beads were finally resuspended in a new tube with 16 μL of 10 mM Tris-HCl at a pH of 8.0.

#### 8. Library preparation and paired-end sequencing

A-tailing of Hi-C DNA was performed by adding 2.5 μL of 10X NEB2 buffer (B7002S), 5 μL of 1 mM dATP (N0440S), and 1.5 μL of Klenow fragment (NEB, M0212L) to the 16 μL of resuspended streptavidin beads. The reaction was incubated for 30 minutes at 37°C in a PCR machine. After the reaction, the beads were settled on a magnetic stand and washed once with 100 μL of BWT buffer and twice with 100 μL of 10 mM Tris-HCl at pH 8.0 (Elution buffer) in new Eppendorf tubes. The streptavidin beads were finally resuspended in 6.5 μL of Elution buffer. Sequencing adaptors were ligated by incubating the beads with 12.5 μL of Quick ligation reaction buffer (NEB, M2200L), 5 μL of ¼ diluted NEBNext Adaptor (NEB, NEBNext Multiplex Oligos for Illumina, E7500S), and 1 μL of Quick T4 ligase (NEB, M2200L). The reaction was incubated at 20°C for 15 minutes. After the incubation, 1.5 μL of USER enzyme mix, part of the NEBNext Multiplex Oligos kit, was added and followed by a 15-minute incubation at 37°C. The beads were then washed three times with 1X BWT buffer, changing tubes at each wash step, and twice with 100 μL of Elution buffer, without changing tubes. Finally, the beads were resuspended in 10 μL of Elution buffer. PCR amplification of the sequencing library was performed by adding 1.25 μL of Universal PCR primer and 1.25 μL of the Index primer of choice, both included in the NEBNext Multiplex Oligos for Illumina kit, as well as 12.5 μL of NEBNext Ultra II Q5 Master Mix (NEB, M0544S). The PCR protocol was as follows: 98°C for 1 minute, followed by 11 cycles of 98°C for 10 seconds, 62°C for 75 seconds, and 72°C for 20 seconds. After cycling, samples were incubated at 65°C for 5 minutes and held at 10°C hold until further processing. PCR products were purified using SPRIselect beads (Beckman Coulter) by adding 20 μL of beads to 25 μL of PCR product (0.8x volume ratio), incubating at room temperature for 15 minutes, washing twice with 80% ethanol, and air-drying the beads. The DNA was finally eluted in 12 μL of Elution buffer, and sequenced using the Illumina NextSeq 500 with 50 bp, paired-end sequencing.

### FISH

#### 1. Oligopaint probe preparation

We used Oligopaint probes and libraries designed in our previous study^23^. Oligopaint probe amplification and purification largely followed the published method^100^. Briefly, the relevant chromosome 1 sub-library was amplified by PCR using sub-library specific primers (synthesized by Eurofins) and 0.5 μL of the corresponding chromosome 1 library pool as DNA template. PCR products were purified using SPRIselect beads (Beckman Coulter) following the manufacturer’s protocol. The purified PCR product was diluted to 20 pg/μL, and 1 μL was used as template for PCR amplification reactions using the common primer pair with the forward primer labeled by Alexa-488 and the 5’ end of the reverse primer phosphorylated (both synthesized by IDT). The second PCR products were pooled together and purified by the addition of 2 μL 20 mg/mL glycogen, 65 μL of 4 M ammonium acetate, and 1350 μL of ice-cold 100% ethanol per 600 μL of PCR reaction. Samples were incubated at -80°C for 35 minutes and centrifuged at maximum speed for 1 hour at 4°C in a tabletop centrifuge. Pellets were subsequently washed with ice-cold 100% ethanol and centrifuged under the same conditions for 30 minutes, air dried at 42°C in a heat block for 15 minutes, and resuspended in 60 μL molecular biology-grade water. Single-strand DNA probes (ssDNA) were generated by treating PCR products with 7.5 μL Lambda exonuclease (NEB, M0262S 10 U/μL) in nicking enzyme buffer for 4 hours at 65°C, followed by heat inactivation at 80°C for 20 minutes. ssDNA was purified following the same ammonium acetate/ethanol precipitation as described above and was resuspended in a final volume of 40 μL molecular biology-grade water. The final concentration yield of ssDNA probe was between 30-40 pmol/μL, and 40 pmol of probe was used per slide for FISH.

#### 2. Oligopaint FISH of YSE cells

Control and lamin-A/B1 DKO YSE cells were isolated, dissociated, fixed, and spun onto glass slides using Cytospin as described above. Cells were briefly washed in PBS, incubated in 0.5% Triton-X-100 for 10 minutes at room temperature in a coplin jar, rinsed in PBS, and incubated overnight in 20% glycerol in a coplin jar. The next day, slides were frozen in liquid nitrogen for 10 seconds and thawed at room temperature; this cycle was repeated 3 times. Slides were next washed with PBS three times for 5 minutes each, incubated in 0.1 M HCl for 5 minutes, and washed with 2X SSC (0.3 M NaCl, 0.03 M sodium citrate) three times for 1 minute each. Slides were then pre-warmed to 37°C for four hours in 2X SSC containing 50% formamide. 40 pmol of ssDNA probes as prepared above were added to the hybridization cocktail (2X SSC, 50% formamide) and was also pre-warmed to 37°C for 2 hours using a heat block. 25 μL of hybridization cocktail containing ssDNA probes was spotted onto a glass coverslip. Slides were inverted onto the coverslip, sealed with rubber cement, denatured at 78°C for 3 minutes using a metal heat block, and hybridized overnight for at least 14 hours in a humidified chamber. The next day, coverslips were removed and slides were washed twice in 50% formamide at 37°C for 30 minutes, once in 20% formamide at room temperature for 10 minutes, followed by four quick washes in 2X SSC. DNA was counterstained with DAPI diluted in 2X SSC for 5 minutes at room temperature. Slides were finally washed three times in 2X SSC for 5 minutes each and mounted using Prolong Gold Antifade. Confocal images were acquired with a Leica SP5 confocal microscope using a 63X objective. Images were acquired as confocal z-stacks with a step size of 0.2 μm using the same laser setting across samples.

#### 3. Quantification of Oligopaint FISH

Imaris software (Bitplane) was used for Oligopaint FISH quantification. To measure the distance from the probe region of chromosome 1 sub-library (Alexa-488) to the nuclear periphery labeled by DAPI, we created ‘Spots’ and ‘Surface’ objects in Imaris. ‘Spots’ objects were created based on Alexa-488 signal from the FISH probe, with manual thresholding applied until the ‘Spots’ objects covered all of the FISH signals. ‘Spots’ objects with a minimum diameter of 1 μm and maximum diameter of 5 μm were retained. ‘Surface’ objects were created based on DAPI staining to outline nuclei. Manual thresholding was applied until most of the Surfaces covered the DAPI-stained nuclei. The distance from the center of each FISH ‘Spot’ to the DAPI-outlined nuclear ‘Surface’ was measured using Imaris’ built-in function. Distance measurements were exported to R for plotting and statistical analysis.

### RNA-seq data analysis

Pre-processing of RNA-sequencing data was performed using Nextflow version 22.10.1 by running the nf-core/rnaseq pipeline, version 3.12.0. Briefly, reads were mapped using the STAR aligner to the mouse genome mm10, and transcript quantification was performed using the Salmon software and GENCODE GTF file (vM23). For differential gene expression analysis, we used the DESeq2 R package. We used the merged gene counts file output from Salmon as input into DESeq2. We removed genes with a total RNA count less than 10 across all replicates from the differential gene expression analysis. We used FDR < 0.05 as a threshold for differential gene expression. Biological process Gene Ontology (GO) term analysis of the differentially expressed genes was performed using the rbioapi R package.

### ATAC-seq data analysis

All ATAC-seq data were processed using the nf-core/atacseq pipeline version 2.1.2, executed with Nextflow version 23.10.0. Reads were aligned to the mm10 mouse reference genome, and the GENCODE GTF file (vM23) annotation was provided during pipeline execution. A read length of 200 was specified using the --read_length flag, which was used as the MACS2 genome size parameter for background estimation during peak calling. The nf-core/atacseq pipeline generated a bed file of broad peaks called by MACS2, which were used for all downstream transcription factor motif analyses. To find ATAC-seq peaks within different LAD categories, we used the bedtools intersect command to overlap the ATAC-seq peaks bed file with merged bed files representing each LAD category. Additionally, we used the annotation of peaks relative to gene features performed by Homer as part of the nf-core/atacseq pipeline to define differentially expressed genes associated with ATAC-seq peaks.

### Transcription factor motif search

All transcription factor motif searches were performed using HOMER version 4.11 software. To find transcription factor motifs enriched in different sets of ATAC-seq peaks, we utilized the <u>findMotifsGenome.pl</u> command with the -size 200 flag, which analyzes a 200 bp window centered around each peak.

### CUT&RUN data analysis

All CUT&RUN data were processed using the nf-core/cutandrun pipeline version 3.2.2, executed with Nextflow version 23.10.0. Reads were aligned to the mm10 mouse reference genome, and mitochondrial reads were removed by including the --remove_mitochondrial_reads flag during pipeline execution.

### LAD mapping CUT&RUN analysis and comparison to LB1-DamID from mESCs

LAD mapping experiments were performed in YSE cells isolated from CD1 strain yolk sacs. For lamin and IgG CUT&RUN experiments, mapped reads from IgG and different lamin isoform CUT&RUN samples were aggregated into 5 kb genomic windows across the mm10 mouse genome using the bedtools coverage command. To reduce background noise, IgG CUT&RUN reads from six biological replicates were merged prior to analysis, as these control experiments yielded fewer reads compared to the lamin CUT&RUN samples. To quantify lamin enrichment across the genome, read numbers at 5 kb resolution from lamin and IgG CUT&RUN samples were normalized using the Counts Per Million (CPM) normalization method. The log₂-fold enrichment was then calculated as the ratio of lamin CPM to IgG CPM for each 5 kb genomic window. These enrichment values were subsequently z-scored and smoothed using a simple moving average (window size = 3 windows). After confirming strong correlation among the lamin-A, lamin-B1, and lamin-B2 CUT&RUN replicates, we averaged the z-scored lamin enrichment values for the three lamin isoforms across 5 kb genomic windows to generate a composite lamin enrichment profile.

To annotate LAD and non-LAD states using our YSE cell lamin CUT&RUN data and published mESC LB1-DamID data, we used the depmixS4 R package to fit a 2-state Gaussian Hidden-Markov model (HMM). To make the mESC LB1-DamID comparable to our YSE LAD mapping, we first used the UCSC liftover tool to convert the mESC LB1-DamID data to mm10 genome coordinates. Next, we used the bedtools map command to calculate the mean LB1-DamID signal across 5 kb genomic windows defined in our YSE LAD mapping data. We next removed genomic regions absent in either dataset from the analysis. To perform HMM calling on our YSE lamin CUT&RUN, we first averaged the log_2_-transformed enrichment values of each lamin isoform (lamin-A, lamin-B1, lamin-B2) across 5 kb genomic windows. These averaged values were then used as input for the HMM, which classified each genomic window into one of two states corresponding to either LADs or non-LADs. We used the same HMM framework to call LADs using log_2_-enrichment values from the published LB1-DamID mESC data. We used the bedtools intersect command with LAD and non-LAD HMM states called in YSE and mESCs to define shared LAD, shared non-LAD, YSE-specific non-LADs, and YSE-specific LAD regions. The resulting bed files for each LAD category were merged using bedtools merge with the flag -d 1.

### Analyses of H3K27ac, H3K27me3, H3K9me2, H3K9me3, and H3K4me3 CUT&RUN from wild-type YSE cells

For analysis of histone post-translational modification CUT&RUN data derived from YSE cells of CD1 embryos, we used the default spike-in normalization method during execution of the nf-core/cutandrun pipeline. H3K4me3 promoter peaks were called during nf-core/cutandrun pipeline execution using MACS2 with default pipeline parameters.

### Analyses of H3K27me3 and H3K27ac CUT&RUN from lamin-A/B1 DKO and control YSE cells

For analysis of H3K27me3 and H3K27ac CUT&RUN data from lamin-A/B1 DKO and control YSE cells, the nf-core/cutandrun pipeline was executed with the --normalisation_mode CPM option to apply counts per million (CPM) normalization. After processing with the nfcore/cutandrun pipeline, bigWig files were generated for each replicate. We used the bigwigMerge command (UCSC Genome Browser tools) to merge replicates belonging to the same genotype, and these aggregate signal profiles were used for downstream analysis. To assess histone mark enrichment, we used the computeMatrix and plotProfile commands from the deepTools package to visualize enrichment of H3K27me3 and H3K27ac at H3K4me3-marked promoters of differentially expressed genes in lamin-A/B1 DKO and control YSE cells.

### HiLands hidden-Markov model analysis

#### 1. Chromatin feature aggregation and preprocessing

We aggregated H3K27me3, H3K27ac, H3K9me2, H3K9me3, and H3K4me3 CUT&RUN data along with ATAC-seq data from wild-type YSE cells into 5 kb windows, corresponding to the same 5 kb windows using in our LAD mapping analyses. We also included the averaged z-scored lamin enrichment values from lamin-A, lamin-B1, and lamin-B2 LAD mapping data for HiLands HMM modeling. Genomic regions corresponding to sex chromosomes were excluded from all analysis. The aggregated CUT&RUN and ATAC-seq data were log2-transformed after adding a pseudocount of 0.001, and outliers were removed from each dataset by capping at the 10th and 90th percentile - any values below the 10th percentile were set equal to the 10th percentile value, and any values greater than the 90th percentile were set equal to the 90th percentile value. We then performed PCA on this dataset consisting of 7 chromatin features using the princomp command in R and kept the first three principal components for further analysis. Uninformative genomic regions with no signal across all chromatin features were removed from the analysis.

#### 2. Hidden-Markov model to assign HiLands

To model HiLands chromatin states in the YSE genome, we applied a HMM framework similar to that described in our previous study^58^. Briefly, we modeled the emission probabilities using a multivariate normal inverse Gaussian distribution and trained the model with a standard expectation-maximization algorithm. To avoid convergence at local minima, the initial parameters of the model were randomized 20 times and the final results were selected based on the highest-likelihood value. Genomic windows were assigned HiLands chromatin states using the Viterbi algorithm. For HiLands HMM modeling in YSE cells, we used a seven-state model. Six chromatin states were found to be informative and had similar characteristics to HiLands previously identified in the mESC genome^58^. The seventh state was uninformative and was found at regions with sparse data, and therefore it was excluded from downstream analyses. All further analyses focused on the six informative HiLands.

#### 3. Comparison of HiLands between YSE cells and mESCs

Previously published HiLands annotations for mESCs were used for all analyses^58^. To analyze LAD transitions between mESC and YSE cells, we focused on mESC HiLands-P and HiLands-B regions within HMM-called mESC LADs and analyzed what HiLands they become in YSE cells. To quantify HiLands state transitions, weighted overlap fractions were calculated from the defined mESC HiLands and corresponding YSE HiLands categories. Regions annotated as uninformative HiLands in YSE cells were excluded from downstream analysis. Transition frequencies were expressed as percentages within each mESC HiLands class.

### Analyses of Hi-C

#### 1. Hi-C data preprocessing

We used the 4DN Hi-C processing pipeline (https://github.com/4dn-dcic/docker-4dn-hic) for read alignment and filtering. Quality control of individual Hi-C libraries was performed independently using the HiCUP analysis pipeline to assess valid read pairs and library complexity. After confirming the high quality of individual Hi-C replicates, deduplicated read pairs from replicates corresponding to each genotype (control or lamin-A/B1 DKO) were merged using the run-merge-pairs.sh script provided in the 4DN pipeline. To generate Hi-C contact matrices, we used the run-juicebox-pre.sh command to make .hic files of the merged control and lamin-A/B1 DKO datasets. Hi-C contact maps were normalized using the Knight-Ruiz (KR) matrix balancing method via the dump command from the Juicer tools suite. Additionally, we generated balanced .cool format Hi-C matrices at 20 kb resolution of the merged control and lamin-A/B1 DKO datasets for downstream TAD inference.

#### 2. TAD Inference

To call TAD boundaries in control and lamin-A/B1 DKO YSE cells, we applied the insulation score method using the insulation function from the cooltools package. Insulation scores were computed from our previously generated 20 kb resolution, balanced .cool Hi-C matrices corresponding to each genotype. For each genomic bin, insulation scores were calculated using a 500 kb sliding window. TAD boundaries were annotated by identifying local minima in the insulation profile. We applied the Li thresholding method^101^ to determine statistically significant insulation drops by including the --threshold Li flag during execution.

#### 3. Hi-C data visualization

For visualization of observed Hi-C reads, we applied a dynamic binning approach as described in our previous study^23^ to the observed contact value matrices in .cool format normalized by total read counts. Dynamic binning was used to reduce noise in low-coverage regions of the Hi-C map. Briefly, each bin pair in the matrix was iteratively expanded in all directions until the total number of contacts within the expanded region met a minimum threshold of 20 reads. The contact frequency for the original bin pair was then calculated as the average of all contact values within this expanded region. This adaptive smoothing ensures that low-signal regions are not disproportionately affected by sampling noise, while preserving resolution in high-signal regions.

We visualized the log_2_-fold change in chromatin interactions between lamin-A/B1 DKO and control using observed/expected (O/E) normalized interactions. This approach highlights differential chromatin interactions between genotypes while controlling for distance-dependent contact decay. Here, we again applied a dynamic binning strategy using observed contact matrices generated by Juicer, which were normalized by the total number of reads to account for differences in sequencing depth. To calculate O/E interaction values for each genotype, we first summed all the observed contact values within a given dynamically expanded bin. We then summed the expected contact values for the same expanded bin. The O/E contact frequency was defined as the ratio of these two sums. To generate expected interaction values for each genotype, we first counted the number of genomic bin pairs at each linear genomic distance that had non-zero interaction values. We then summed the total observed interactions for genomic bins separated by a given linear distance. Lastly, we divided the total observed interactions at each linear distance by the corresponding number of valid window pairs separated by that same linear distance. Finally, these expected values were normalized to sum to one across all distances. This process was conducted independently for each genotype, ensuring that control and lamin-A/B1 DKO datasets had their own matched expected interaction profiles for accurate and unbiased downstream analysis.

#### 4. Hi-C interaction change analysis between different HiLands

To assess global intra-chromosomal interaction changes between different HiLands, we calculated the log₂ fold-change in chromatin interactions between lamin-A/B1 DKO and control cells using observed/expected (O/E) normalized interaction frequencies, following a similar approach as described above. For this analysis, we used observed contact matrices generated by Juicer at 5 kb resolution, which were normalized by the total number of reads. We first calculated the total number of interacting window pairs at each linear genomic distance for all the pairwise combinations of HiLands. To compute the O/E interaction ratio for each HiLands pair, we first summed all observed interaction values genome-wide for that specific HiLands pair. Then, for each genomic distance, we multiplied the number of window pairs by the expected contact frequency at that distance and subsequently summed all those values to obtain the denominator of the O/E ratio for each HiLands pair. The O/E interaction ratio for each HiLands pair was calculated by dividing the total observed interactions by the summed expected values calculated above. Finally, we computed the log₂ fold-change of lamin-A/B1 DKO O/E interactions over control O/E interactions for each pair of HiLands to quantify changes in chromatin interaction patterns genome-wide upon lamin-A/B1 loss. An analogous approach was used to calculate changes in interactions between promoters of differentially expressed genes and the various HiLands upon lamin-A/B1 loss. Interaction changes were assessed with HiLands regions located within 5 million base pairs upstream and downstream of a given gene promoter. For heatmap visualization, values were capped at the 25^th^ and 75^th^ percentile to reduce the influence of extreme values and improve visual interpretability. To assess the statistical significance of Hi-C interaction changes for upregulated and downregulated genes, we compared their promoter interaction changes with each HiLand category to the promoter interaction changes observed for a randomized control set of genes. Each control set of genes was selected from the same LAD category region and included genes with Salmon RNA-seq counts greater than 10. Differentially expressed genes that we identified by DESeq2 (FDR < 0.05) were excluded from the control sets of genes.

#### 5. Hi-C compartmentalization analysis

We calculated A/B compartment scores using the C-score tool previously developed in our lab^64^. Briefly, compartment values were computed at 20 kb resolution using the raw deduplicated pairs file output generated for each genotype from the 4DN pipeline. A minimum interaction distance of 1 Mb was used to exclude short-range contacts from the calculation, which are typically dominated by TAD-level interactions. After C-score inference, we compared the C-scores to our LAD mapping data to guide compartment annotation. C-scores for a given chromosome were multiplied by -1 if necessary to ensure that positive C-scores corresponded to the B compartment as defined by our CUT&RUN LAD mapping. To visualize compartmentalization patterns in control and lamin-A/B1 DKO YSE cells, C-scores for each genomic bin were smoothed using a three-bin simple moving average.

## QUANTIFICATION AND STATISTICAL ANALYSES

Quantification methods, including the number of biological replicates, sample size, and statistical tests used, can be found in the figure legends. Sample sizes were not determined by a priori power calculations, and underlying data distributions were not tested for normality. Samples were not randomized. Significance was defined as *p* < 0.05 unless otherwise noted.

## Supplementary Figure Legends

**Figure S1. Related to Figure 1:**
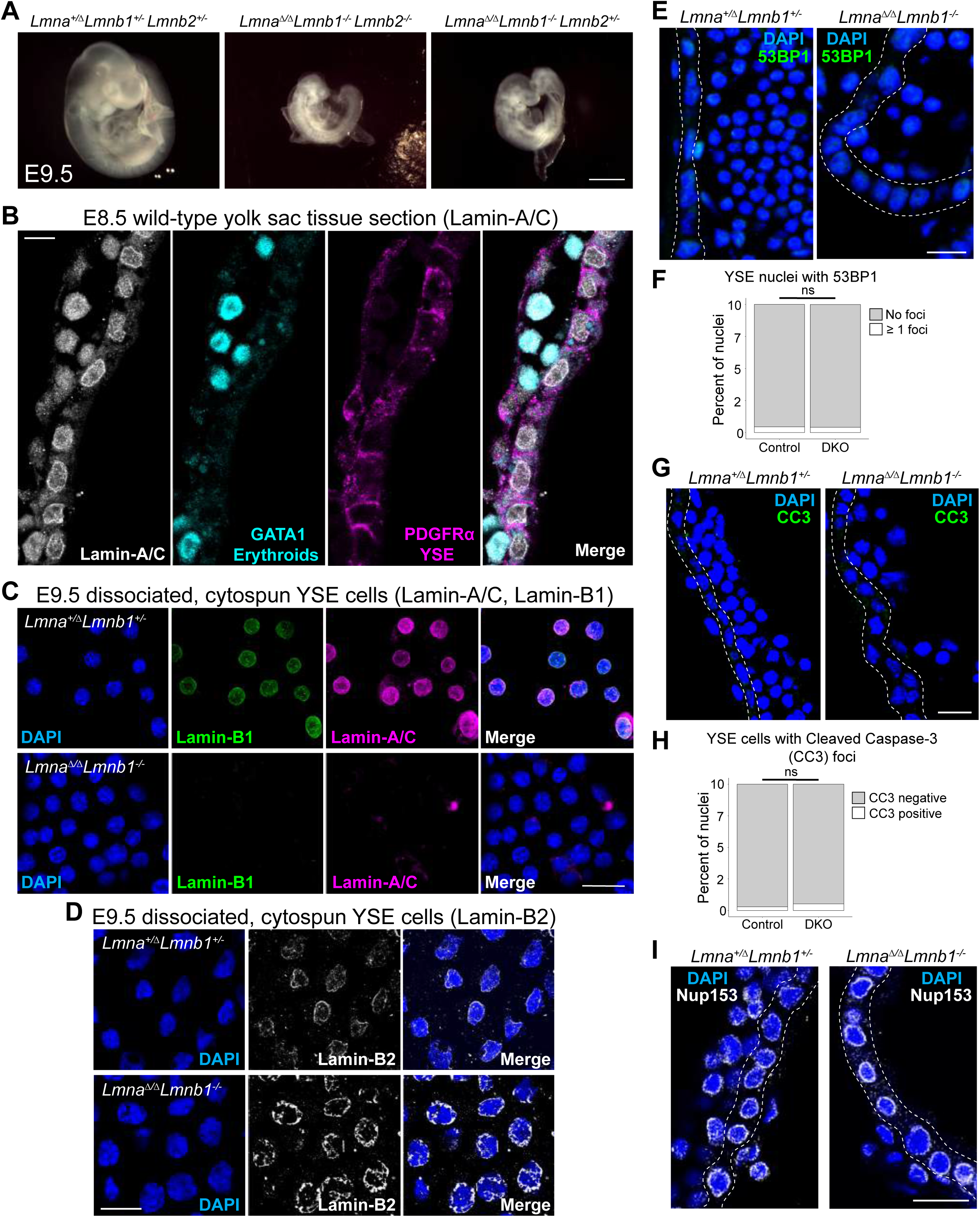
Further characterization of lamin triple-knockout embryos and lamin-A/B1 double-knockout (DKO) yolk sac endoderm (YSE) cells. (A) Representative images of *Lmna^+/Δ^Lmnb1^+/-^Lmnb2^+/-^*, *Lmna^Δ/Δ^Lmnb1^-/-^Lmnb2^-/-^*, and *Lmna^Δ/Δ^Lmnb1^-/-^Lmnb2^+/-^* embryos at embryonic day (E)9.5. Scale bar, 500 µm. (B) Representative lamin-A immunofluorescence staining in a E8.5 wild-type yolk sac tissue section. Red blood cells are marked by GATA1 immunofluorescence, and YSE cells are marked by PDGFRα. Nuclear peripheral lamin-A staining is evident in YSE cells. Scale bar, 10 µm. (C) Representative images of lamin-A and lamin-B1 immunofluorescence staining of dissociated control and lamin-A/B1 DKO YSE cells cytospun onto glass slides. Lamin-A and lamin-B1 staining is absent in lamin-A/B1 DKO YSE cells. The residual signal observed upon lamin-A staining of lamin-A/B1 DKO YSE cells corresponds to non-specific background due to reactivity of the anti-mouse secondary antibody with endogenous IgG present in YSE cells. Scale bar, 20 µm. (D) Representative images of lamin-B2 immunofluorescence staining in dissociated and cytospun control and lamin-A/B1 DKO YSE cells. Scale bar, 20 µm. (E) Representative images of immunofluorescence staining for 53BP1, a DNA damage marker, in control and lamin-A/B1 DKO YSE cells from yolk sac tissue sections. Scale bar, 20 µm. (F) Bar plot showing the percentage of control and lamin-A/B1 DKO YSE cells exhibiting 53BP1 foci. N = 2 biological replicates; 255 control nuclei and 269 lamin-A/B1 DKO nuclei analyzed. Fisher’s exact test Odds ratio: 1.18, *p* = 0.83 (not significant - ns). (G) Representative images of immunofluorescence staining for cleaved caspase-3 (CC3), an apoptosis marker, in control and lamin-A/B1 DKO YSE cells from yolk sac tissue sections. Scale bar, 20 µm. (H) Bar plot showing the percentage of control and lamin-A/B1 DKO YSE cells exhibiting CC3 foci. N = 2 biological replicates, 171 control nuclei and 169 lamin-A/B1 DKO nuclei analyzed. Fisher’s exact test Odds ratio: 0.535, *p* = 0.29 (not significant - ns). (I) Representative images of immunofluorescence staining for NUP153, a component of the nuclear pore complex, in control and lamin-A/B1 DKO YSE cells from yolk sac tissue sections. Scale bar, 20 µm.

**Figure S2. Related to Figure 2:**
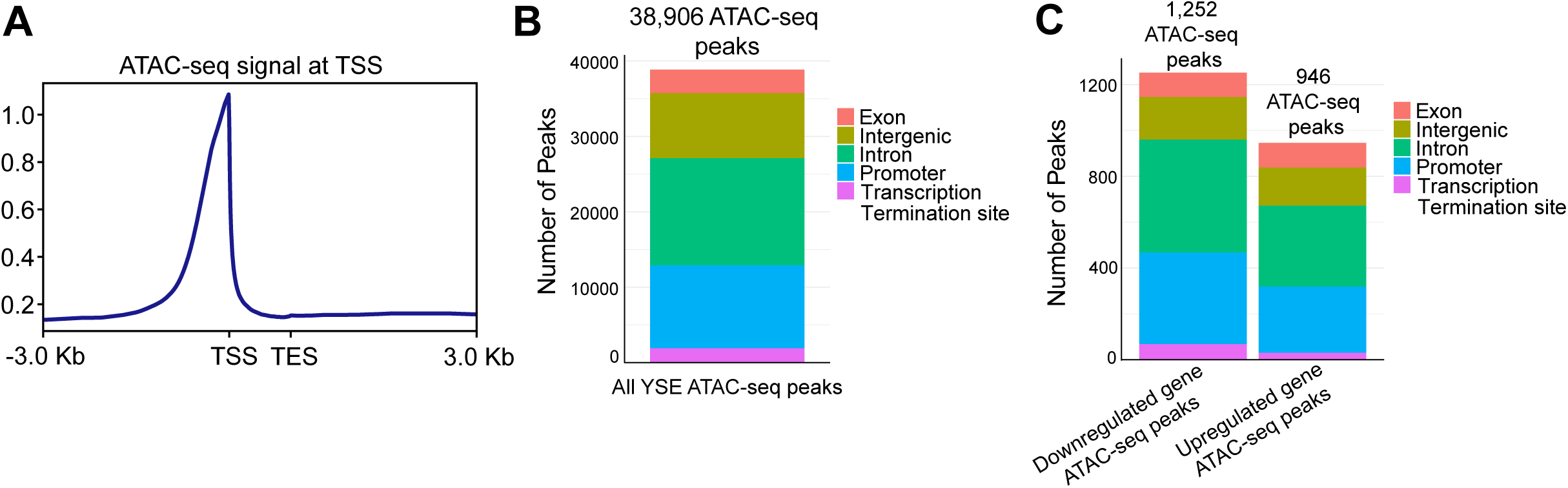
ATAC-seq and peak calling quality control. (A) Metaplot showing ATAC-seq signal enrichment around transcription start sites (TSSs) in YSE cells. (B) Grouped bar chart showing the number of ATAC-seq peaks identified in YSE cells, color-coded by their distribution across exons, intergenic regions, introns, promoters, and transcription termination sites. (C) Grouped bar chart showing the number of ATAC-seq peaks associated with genes downregulated or upregulated upon lamin-A/B1 loss in YSE cells. Bars are color-coded by the distribution of peaks across exons, intergenic regions, introns, promoters, and transcription termination sites.

**Figure S3. Related to Figure 3:**
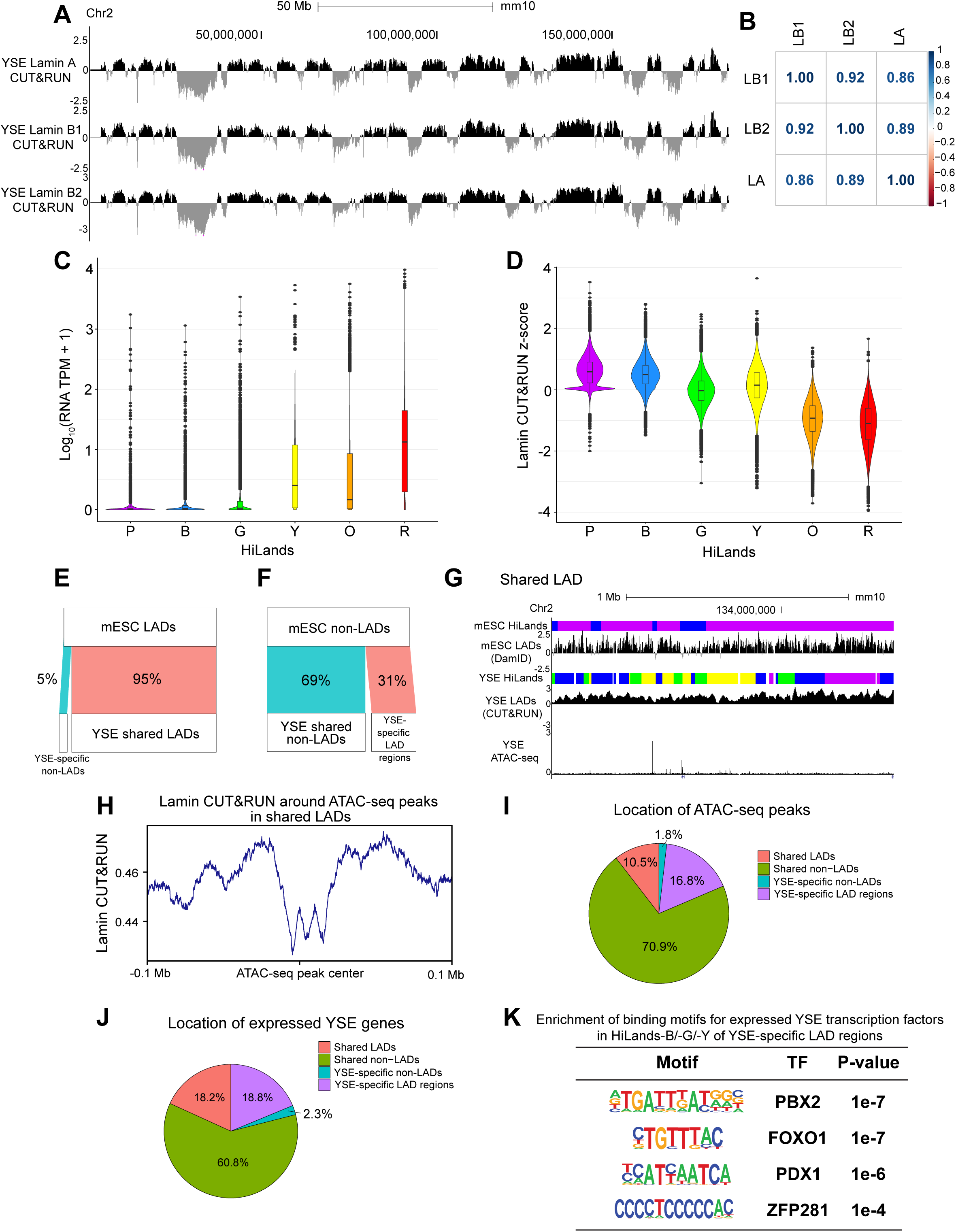
Lamin CUT&RUN quality control and additional characterization of HiLands and LAD remodeling in YSE cells. (A) Genome browser view of chromosome 2 showing CUT&RUN profiles generated using antibodies against the three lamin isoforms lamin-A, lamin-B1, and lamin-B2. Tracks are visualized as lamin CUT&RUN enrichment over IgG CUT&RUN. (B) Pearson correlation coefficients between the z-scored CUT&RUN enrichment profiles of each lamin isoform normalized to IgG CUT&RUN. (C) Violin plots with overlaid box plots showing the distribution of wild-type YSE gene expression levels based on bulk RNA-seq across the six YSE HiLands (TPM – transcripts per million). Boxes show the interquartile range and center lines indicate medians. (D) Violin plots with overlaid box plots showing the distribution of z-scored lamin CUT&RUN enrichment across the six YSE HiLands. Boxes show the interquartile range and center lines indicate medians. (E and F) Alluvial plots showing the transition of mESC LADs to corresponding LAD or non-LAD regions in YSE cells (E) and the transition of mESC non-LADs to corresponding LAD or non-LAD regions in YSE cells (F). LADs and non-LADs were called using a 2-state hidden-Markov model. (G) Example of a shared LAD between YSE cells and mESCs. Genome browser tracks show published mESC LADs mapped by lamin-B1 Dam-ID and mESC HiLands, and the corresponding YSE HiLands, YSE LADs mapped by lamin CUT&RUN, and YSE ATAC-seq peaks generated in this study. (H) Metaplot showing lamin CUT&RUN enrichment around YSE ATAC-seq peaks associated with the shared LAD regions. (I) Pie chart showing the distribution of ATAC-seq peaks in YSE cells across genomic regions corresponding to the four categories of LADs and non-LADs. (J) Pie chart showing the distribution of expressed genes in YSE cells across genomic regions corresponding to the four categories of LADs and non-LADs. (K) Enrichment of binding motifs for transcription factors expressed in YSE cells in YSE-specific LAD regions characterized by HiLands-B, -G, and -Y.

**Figure S4. Related to Figure 4:**
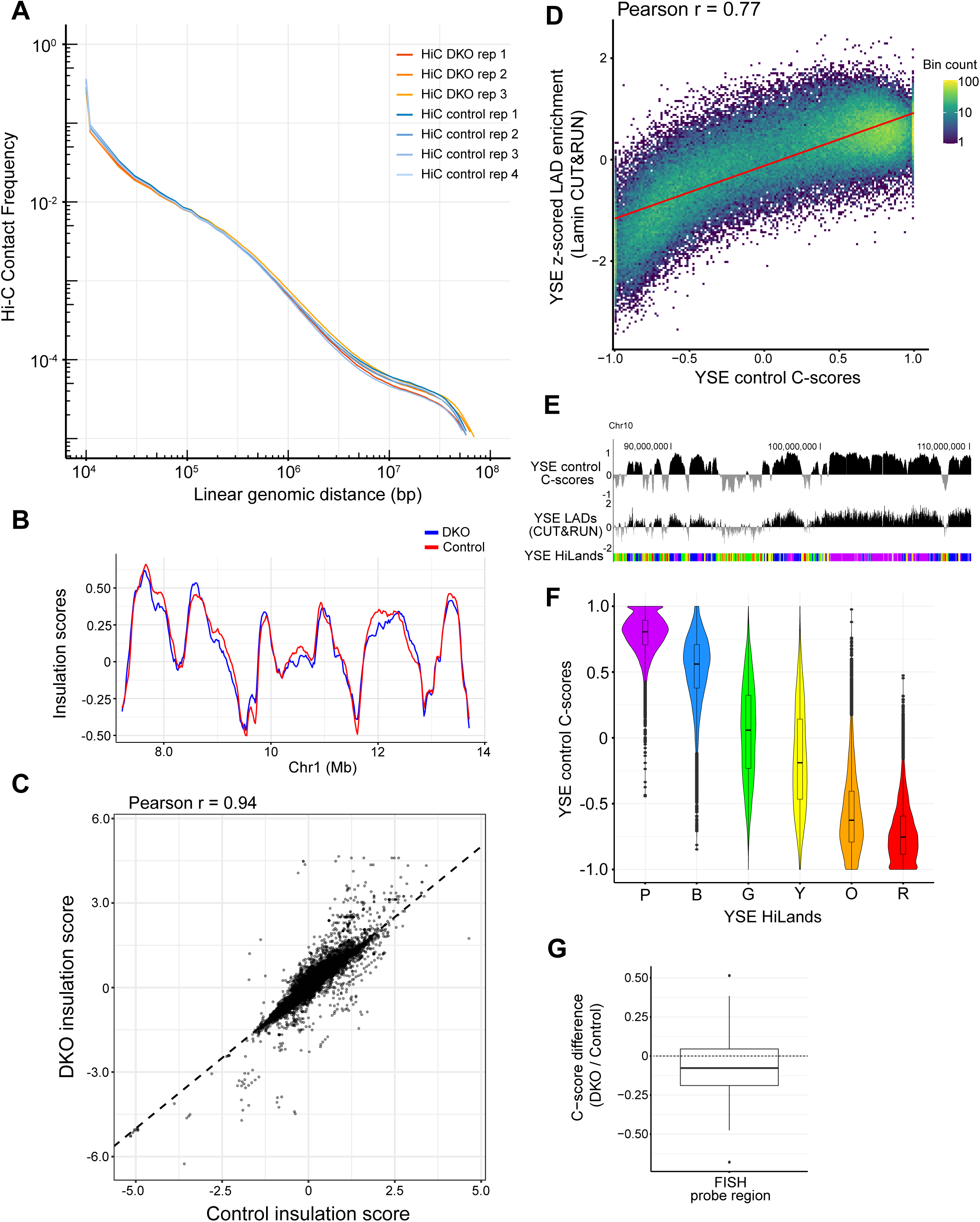
Hi-C quality control and additional characterization of TAD boundaries and compartment scores (C-scores). (A) Hi-C contact decay curves showing the decrease in Hi-C contact frequency as a function of linear genomic distance for individual Hi-C replicates from control and lamin-A/B1 DKO YSE cells. (B) Insulation scores calculated from control and lamin-A/B1 DKO YSE cells for the representative chromosome 1 region shown in Figure 4A and 4B. (C) Genome-wide scatterplot comparing insulation scores from control and lamin-A/B1 DKO YSE cells. The dashed line indicates line of best fit. Pearson correlation coefficient, r = 0.94. (D) Density plot comparing z-scored enrichment of wild-type YSE LADs determined by lamin CUT&RUN to C-scores calculated from control YSE Hi-C data. The red line indicates line of best fit. Pearson correlation, r = 0.77. (E) Representative genome browser tracks showing C-scores from control YSE cells alongside YSE LADs mapped by lamin CUT&RUN and YSE HiLands. (F) Violin plots with overlaid box plots showing the distribution of control C-scores across the six YSE HiLands. Boxes show the interquartile range and center lines indicate medians. (G) Box plot showing decreased C-scores upon lamin-A/B1 loss within the FISH probed region. Boxes show the interquartile range, center lines indicate medians, and whiskers extend to 1.5 times the interquartile range.

**Figure S5. Related to Figure 5:**
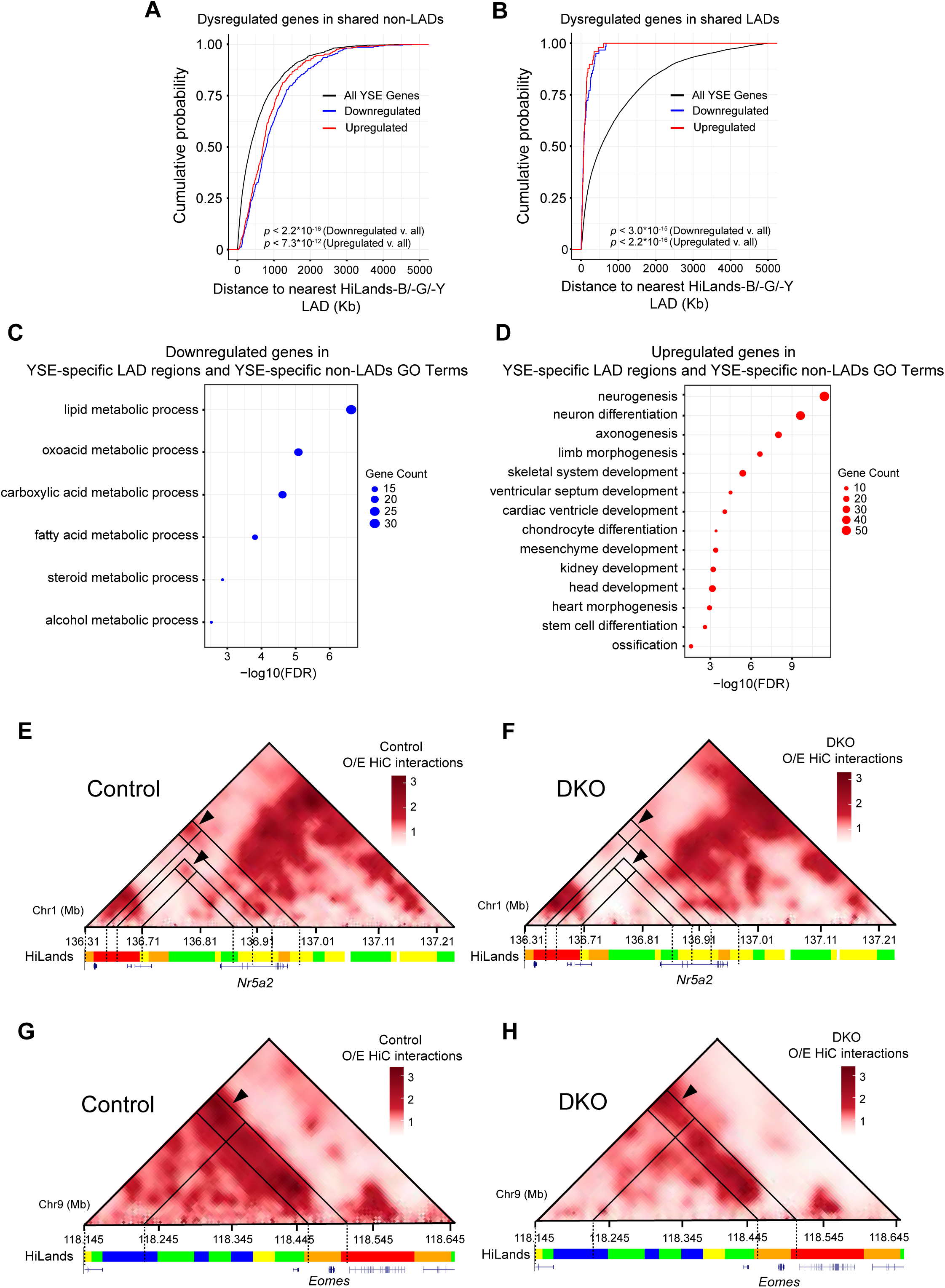
Additional characterization of 3D chromatin interaction changes around dysregulated genes in YSE-specific LAD regions and YSE-specific non-LADs. (A and B) Cumulative probability plot of distances from dysregulated gene promoters in shared non-LADs (A) or shared LADs (B) to the nearest LADs characterized by HiLands-B, -G, and -Y. Promoter distances of all expressed YSE genes are shown as a control. (C and D) Selected GO terms significantly enriched among differentially expressed genes located in YSE-specific LAD regions and YSE-specific non-LADs that are downregulated (C) or upregulated (D) following lamin-A/B1 loss. (E and F) Heatmaps of observed/expected (O/E) Hi-C interactions in control (E) and lamin-A/B1 DKO (F) YSE cells around the downregulated gene *Nr5a2*. Boxed regions and arrows highlight decreased chromatin interactions between *Nr5a2* and transcriptionally active HiLands-R upon lamin-A/B1 loss. (G and H) Heatmaps of observed/expected (O/E) Hi-C interactions in control (G) and lamin-A/B1 DKO (H) YSE cells around the upregulated gene *Eomes*. Boxed regions and arrows highlight decreased chromatin interactions between *Eomes* and repressive HiLands-B upon lamin-A/B1 loss.

**Figure S6. Related to Figure 6:**
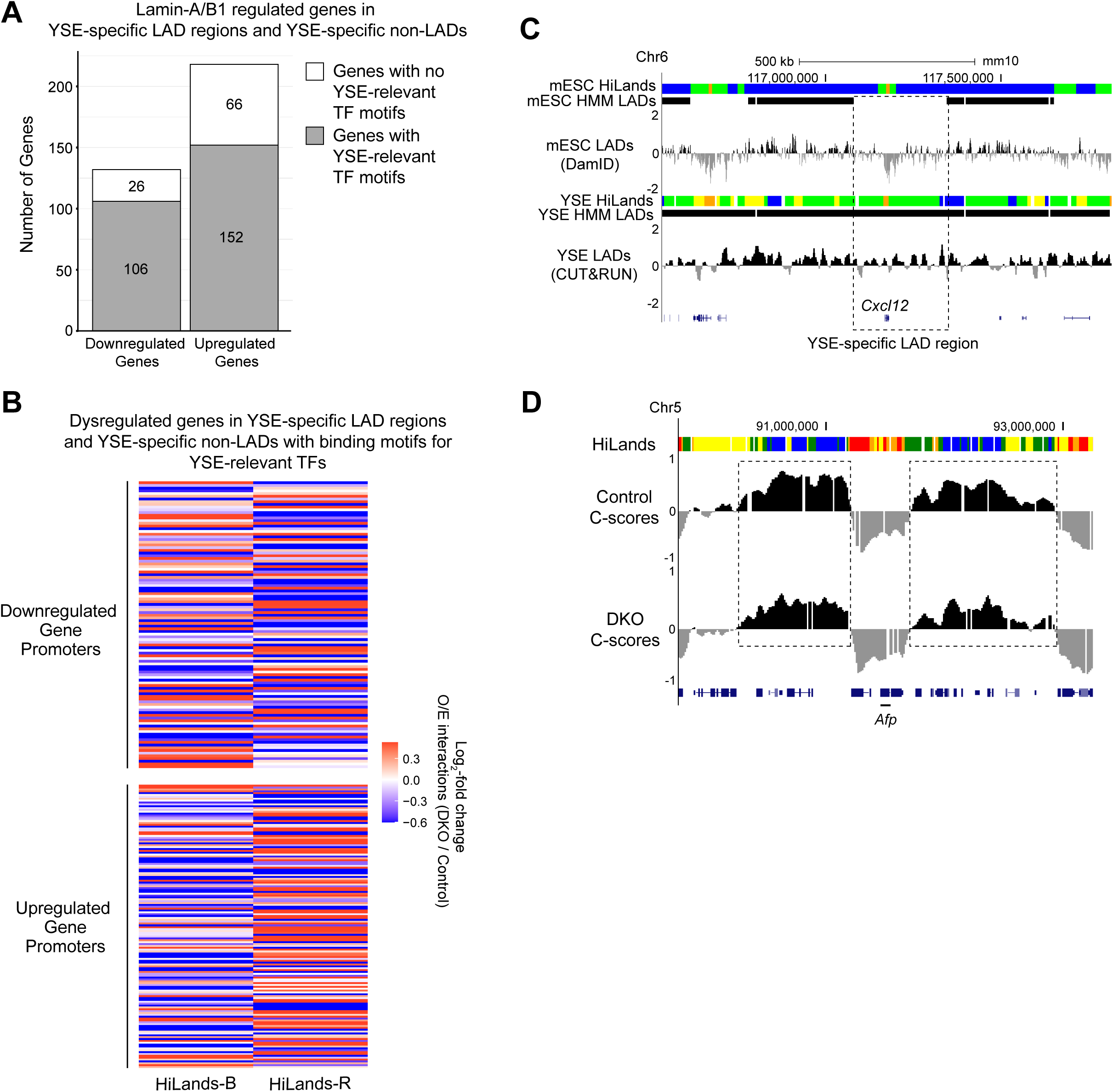
Additional characterization of dysregulated genes upon lamin-A/B1 loss containing YSE-relevant transcription factor motifs. (A) Bar chart showing the number of dysregulated genes located in YSE-specific LAD regions and YSE-specific non-LADs that contain binding motifs of YSE-relevant transcription factors. (B) Heatmaps showing interaction changes of 5 kb bins surrounding promoters of dysregulated genes associated with binding motifs of YSE-relevant transcription factors in the indicated LAD and non-LAD regions. Interaction changes with HiLands-B and HiLands-R regions within ±5 Mb of each promoter are shown. Downregulated genes exhibit decreased interactions with HiLands-R (*p* = 0.043, Wilcoxon rank-sum test), while upregulated genes display decreased interactions with HiLands-B (p = 0.013, Wilcoxon rank-sum test). Significance was assessed relative to randomized control genes. (C) Genome browser tracks showing the upregulated broad-lineage gene *Cxcl12* alongside published mESC LADs mapped by lamin-B1 DamID, published mESC HiLands, and the mESC HMM-called LAD annotations generated in this study, as well as the YSE HMM-called LAD annotations, YSE HiLands, and YSE LADs mapped by lamin CUT&RUN in this study. The dashed box indicates the YSE-specific LAD region encompassing *Cxcl12*. (D) Genome browser view showing Hi-C compartment scores (C-scores) from control and lamin-A/B1 DKO YSE cells alongside YSE HiLands surrounding the *Afp* locus. Dashed boxes indicate HiLands-B, -G, and -Y regions exhibiting decreased C-scores upon lamin-A/B1 loss in YSE cells.

## Supplemental Tables

**<u>Table S1.</u>** List of significantly downregulated (Sheet 1) and significantly upregulated (Sheet 3) genes upon lamin-A/B1 loss based on bulk RNA-seq analysis of control and lamin-A/B1 DKO E9.5 YSE cells. Gene Ontology (GO) term analysis of significantly downregulated (Sheet 2) and significantly upregulated (Sheet 4) genes upon lamin-A/B1 loss in YSE cells, Related to Figure 1.

**<u>Table S2.</u>** Motif analyses of ATAC-seq peaks associated with significantly downregulated (Sheet 1) and significantly upregulated (Sheet 2) genes, Related to Figure 2.

**<u>Table S3</u>.** Motif analyses of ATAC-seq peaks associated with shared LADs (Sheet 1), YSE-specific non-LADs (Sheet 2), YSE-specific LAD regions (Sheet 3), and motif analyses of HiLands-B, -G, and -Y regions within YSE-specific LAD regions (Sheet 4), Related to Figure 3.

**<u>Table S4.</u>** Quality control statistics of control and lamin-A/B1 DKO Hi-C libraries, Related to Figure 4.

**<u>Table S5.</u>** GO term analysis of differentially expressed genes associated with YSE-specific LAD regions and YSE-specific non-LADs (Sheet 1-2), whose expression changes correlate with 3D chromatin interaction changes. Identification of upstream regulators in this gene set, including Nr5a2 and Eomes (Sheet 3). Analysis of differentially expressed genes characterized by ATAC-seq peaks with *Nr5a2* motifs (Sheet 4) or *Eomes* motifs (Sheet 5), Related to Figure 5.

**<u>Table S6.</u>** Characterization of binding motifs for YSE-relevant transcription factors associated with differentially expressed genes in YSE-specific LAD regions and YSE-specific non-LADs. Related to Figure 6.

**<u>Table S7.</u>** List of primers used for embryo genotyping, Related to STAR Methods.

